# Modelling mucus clearance in sinuses: thin-film flow inside a fluid-producing cavity lined with an active surface

**DOI:** 10.1101/2024.09.08.611783

**Authors:** Nikhil Desai, Eric Lauga

## Abstract

The paranasal sinuses are a group of hollow spaces within the human skull, surrounding the nose. They are lined with an epithelium that contains mucus-producing cells and tiny hairlike active appendages called cilia. The cilia beat constantly to sweep mucus out of the sinus into the nasal cavity, thus maintaining a clean mucus layer within the sinuses. This process, called mucociliary clearance, is essential for a healthy nasal environment and disruption in mucus clearance leads to diseases such as chronic rhinosinusitis, specifically in the maxillary sinuses, which are the largest of the paranasal sinuses. We present here a continuum mathematical model of mucociliary clearance inside the human maxillary sinus. Using a combination of analysis and computations, we study the flow of a thin fluid film inside a fluid-producing cavity lined with an active surface: fluid is continuously produced by a wall-normal flux in the cavity and then is swept out, against gravity, due to an effective tangential flow induced by the cilia. We show that a steady layer of mucus develops over the cavity surface only when the rate of ciliary clearance exceeds a threshold, which itself depends on the rate of mucus production. We then use a scaling analysis, which highlights the competition between gravitational retention and cilia-driven drainage of mucus, to rationalise our computational results. We discuss the biological relevance of our findings, noting that measurements of mucus production and clearance rates in healthy sinuses fall within our predicted regime of steady-state mucus layer development.

## 1 Introduction

The human skull contains air-filled cavities around the nose region, called paranasal sinuses. These are named after the bones in the skull within which they reside: frontal, maxillary, ethmoid and sphenoid [1] (see illustration in Fig. 1a). The sinuses are believed to serve a number of evolutionary and functional purposes, including keeping the skull light and buoyant, imparting resonance to voice, humidifying inspired air, improving olfaction, absorbing physical trauma and producing mucus [2, 3].

**Fig. 1:**
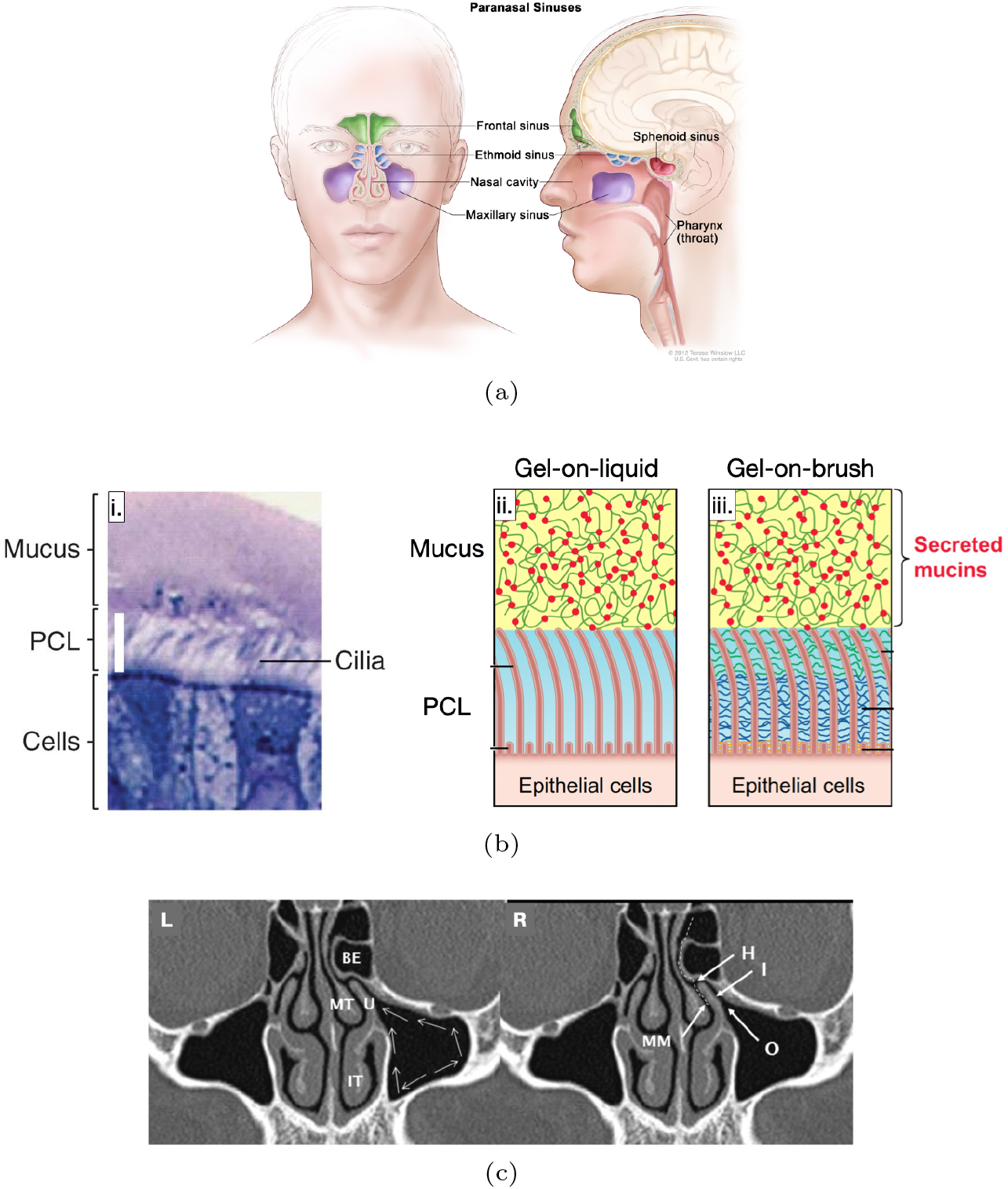
Mucociliary clearance in human sinuses. (a) Sketch of sinuses and their locations inside the human skull [16]. (bi): light microscopy image of the nasal epithelium, showing goblet cells, cilia, the periciliary layer (PCL) and the mucus layer (ML) (reproduced with permission from Ref. [10]). (bii): sketch showing the position of the cilia tips and the interface between the PCL and the ML, for the gel-on-liquid model. (biii): sketch showing the interface between the PCL and the ML, for the gel-on-brush model (reproduced with permission from Ref. [6]). (c) The maxillary sinuses as seen on a CT scan of a human head. The expected direction of mucus flow due to ciliary beating is shown via the thin white arrows in the left (**L**) sub-panel. The sinus exit, called the ostium, is marked by the letter “O” in the right (**R**) sub-panel (reproduced with permission from Ref. [17]).

One role of sinuses is to supply mucus to the nasal cavity, where it plays an important role in the respiratory system [4]. The sinus interior is lined with an epithelium which contains two types of cells: (i) goblet cells that secrete gel-forming proteins called mucin, and, (ii) ciliated cells endowed with active hairlike appendages called cilia [5]. The epithelium is hydrated through osmosis regulated by ion transport across the epithelial cells [6]. The secreted mucins expand drastically upon contacting the hydrated epithelium and form a gel-like fluid called mucus [7]. In this way, mucus is effectively produced inside the sinuses through a combination of mucin-secretion by goblet cells and osmosis-induced hydration of the epithelium. The typical composition of mucus is: ≈ 97.5% water, ≈ 0.5% mucin proteins, ≈ 1.1% each of salts and ≈ 0.9% other globular proteins [6]. It is a bi-layered viscoelastic fluid consisting of a highly viscous mucus layer (ML) overlying a periciliary layer (PCL) which itself rests atop the nasal epithelium [8, 9]. The PCL has been classically postulated to be a water-like fluid layer, but more recent investigations have questioned this gel-on-liquid description, instead proposing that the PCL has a brush-like structure owing to various secreted mucins and other polymers adhered to the epithelium [10]. Regardless, the cilia are immersed almost entirely in the PCL with only their tips penetrating into the ML [11, 12] (see Fig. 1b). They perform coordinated motion in a forward and recovery stroke such that they push the mucus during the forward stroke but cause minimal backflow during the recovery stroke; thus, on average, the mucus blanket is transported along the sinus epithelium [12, 13]. The net effect of cilia-induced mucus transport is that the mucus exits the sinus through an opening called the ostium, and is then directed into the nasopharynx [14] (see Fig. 1c). In this way, a fresh mucus layer is always maintained inside a healthy sinus: being produced continuously at the epithelium, and being cleared out simultaneously by the beating cilia. This process, of constant production and replenishment of mucus, is referred to as mucociliary clearance (MCC) [15].

MCC is a robust process that is responsible for health and defence of the nose, for example, almost all of the particulate matter of size *<* 10 *µ*m that we breathe gets trapped in the mucus and removed before it can cause harm to the underlying tissue [4]. Importantly, inhaled bacteria are removed by MCC before they get time to replicate and become infectious. Any impairment in MCC can cause mucus build-up inside the sinuses; conversely, situations causing excess mucus production (e.g., allergen-induced inflammation of the sinonasal mucosa) can impair MCC and lead to further mucus build-up in the sinuses. These malfunctions are conducive for bacteria to colonise the sinuses, leading to the development of bacterial biofilms and subsequent diseases such as chronic rhinosinusitis [18]. It is therefore crucial to understand the physical factors affecting mucus flow in the sinuses, particularly in the maxillary sinus, which is the main site for sinus disease [19].

A number of interesting fluid flow phenomena govern the evolution and transport of mucus inside the maxillary sinus. Firstly, in order to maintain a steady mucus layer over the sinus walls, there must exist a balance between the rates of mucus production and mucus expulsion due to transport facilitated by ciliated cells. In fact, the thickness of the mucus layer–itself an indicator of susceptibility to disease [20]–would depend on the relative rates of mucus production and mucus clearance. Secondly, since the maxillary sinus ostium is located above the bottom side of the sinus [17], the flowing mucus must overcome gravity in order to successfully exit the sinus (see Fig. 1c). Indeed, it has been clinically postulated that the proclivity of the maxillary sinus to infections is likely due to its ostium being located against the direction of gravity [21– 23]. Thirdly, the mucus layer inside the sinus is exposed to air and can deform due to surface tension, which can then affect its flow.

In this paper, we employ fundamental concepts from fluid mechanics to understand how the aforementioned physical effects interact with each other and contribute to maintain a thin mucus layer inside the maxillary sinus. We first propose in Section 2 a model system that includes relevant bio-physical components dictating mucus flow inside the sinus. The system is comprised of a cavity lined with a fluid-producing active surface, i.e. the inner surface of the cavity produces mucus, and also drives it along the cavity with a prescribed tangential velocity which models the mean action of the cilia on the mucus. In Section 3, we derive a nonlinear evolution equation for the thickness of the mucus film, based on important modifications to classical theories on thin-film flow [24–27]; this is done for both two-dimensional and three-dimensional cavities. In Section 4, we solve this equation numerically to study the nature of mucus film profiles inside the model sinus. Specifically, we determine a phase space, defined by the rates of mucus production and clearance, consisting of two types of solutions: unsteady solutions corresponding to physical conditions that don’t result in successful MCC, and steady solutions for physical conditions that do result in successful MCC from the sinus. We rationalise this demarcation between the unsteady and steady solutions using a physical argument resulting in a scaling relationship in Section 5. We show that, for a prescribed rate of mucus production, successful MCC is achieved only if the cilia-induced mucus flow exceeds a certain threshold; in the process, we identify how this threshold clearance rate scales with the rate of mucus production. In Section 6, we next discuss the direct biological application of our findings by comparing our predictions of steady-state conditions in the model sinus (i.e. rates of mucus production and clearance) to the existing literature on these operating conditions in healthy sinuses. We finish by a summary of our work in Section 7 along with suggestions for future investigations.

## 2 Mathematical model

### 2.1 Key biophysical ingredients

What are the essential ingredients for a minimal model of MCC in the maxillary sinuses? Firstly, it must consist of a finite-size cavity with an outlet for the fluid (mucus) to exit. Biologically, these represent, respectively, the sinus and the ostium (the small opening in the sinus that drains into the nasal cavity). Secondly, there must be some mechanism for fluid production inside the cavity, to model the continuous production of mucus in the sinus. Thirdly, there must also be an active mechanism to continually drive the produced fluid out, modelling the action of the ciliated cells inside the sinus. In healthy conditions, there exists a steady mucus layer in the sinus, which is continuously replenished on account of a balance between mucus production and mucus clearance. This fundamental feature should emerge in our model as a consequence of the forces governing fluid motion.

### 2.2 The minimal model: simplifying assumptions

Based on this, we can propose a simple model, which includes all the above-mentioned biophysical effects. For the sinus cavity, we consider two elementary geometries: a circle or a sphere. The former will be used for a planar/two-dimensional analysis whereas the latter for an axisymmetric/three-dimensional analysis. We treat the mucus, in this first exploration, as a Newtonian fluid with uniform physical properties (viscosity and density). We assume that the mucus is continually produced at the walls of the cavity and that it enters the cavity normally (i.e. perpendicular to the local cavity wall) at a constant velocity 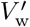 (red arrows in Fig. 2). Without ciliary function, gravity (green arrows in Fig. 2) would cause the mucus to accumulate inside the cavity and fill it up. But cilia actively sweep the mucus up along the wall and cause it to exit the cavity; the effective action of the cilia is thus modelled as an active (or ‘slip’) tangential velocity of characteristic magnitude 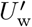, prescribed along the walls of the cavity (blue arrows in Fig. 2). The mucus exits the system at the top through an ostium which is modelled differently in the two geometries. For the circular geometry, we model the mucus exit as a discontinuity: once the mucus reaches the top-most point (orange dot near the top in Fig. 2(a)) it is removed from the domain. For the spherical geometry, we truncate the sphere near its top pole to form a small circular opening from where the mucus exits the domain (orange circle near the top in Fig. 2(b)). We will see that this minimal model is sufficient to explain the development of a thin mucus film inside the sinus (Section 6), and will revisit the various assumptions behind the model when offering perspectives for future work (Section 7.3).

**Fig. 2:**
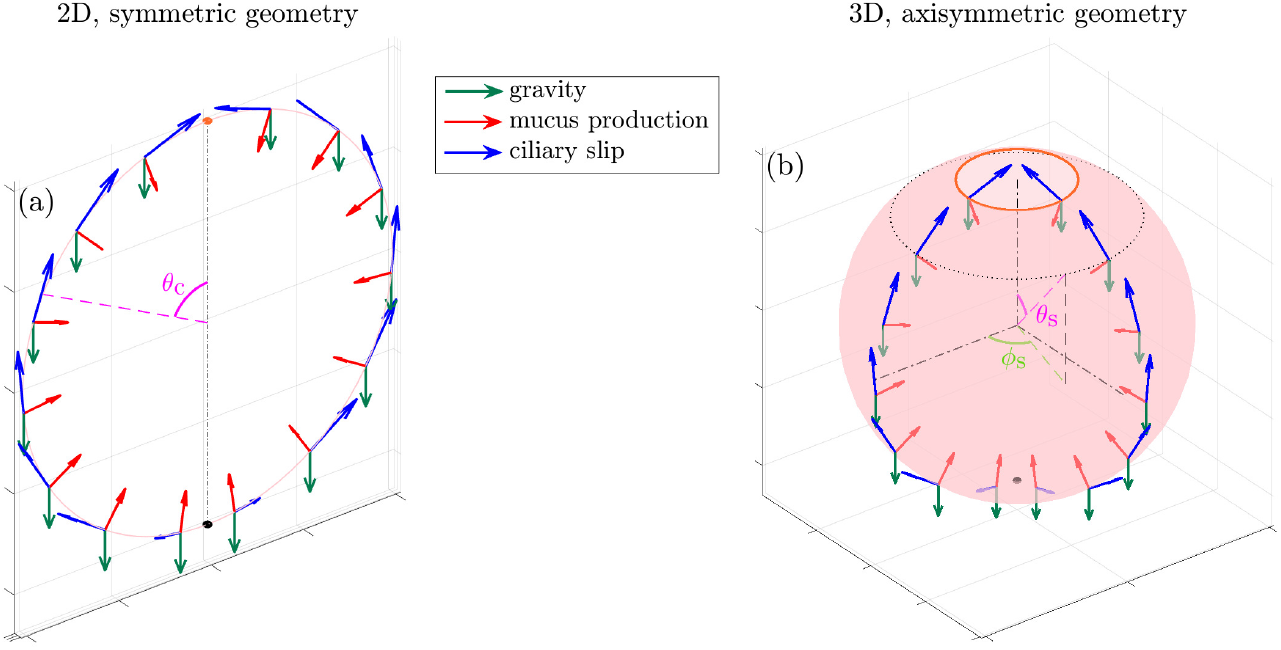
Schematics explaining the geometry of the model sinus, (a) a two-dimensional, circular system, and, (b) a three-dimensional, but axisymmetric spherical system. The blue arrows denote the direction of the effective ciliary slip velocity 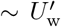, the red arrows denote the wall-normal mucus in-flow 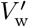, and the downward pointing green arrows denote the direction of gravity. The bottom-most point in both the cases–from where begins the upward motion of the mucus due to cilia action–is marked by a black dot. The mucus exits the system as soon as it reaches the top: (a) the orange dot in the 2D case, and, (b) the orange circle in the 3D case. In panel (b) the velocity vectors are shown for only two azimuths, for clarity, but they are distributed axisymmetrically– around the vertical axis–over the entire sphere surface.

### 2.3 Biologically relevant parameter values

We summarise in Table 1 the values of the various important parameters involved in the problem; note that the physical properties of the mucus, especially its effective viscosity and surface tension, can vary over a range of magnitudes, depending on the general health of the nose [28–30]. In humans, the mucus develops over the sinus epithelium as a film of thickness *h*^′^ ∼ 10-15 *µ*m [14]. The coordinated beating of cilia moves this mucus layer at an average rate of 2-25 mm/min [4, 14, 17], which means that 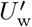 lies in the (large) range 30 to 400 *µ*m/s. To estimate typical values of the mucus production 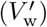 under steady operative conditions, we use a mass balance argument along with measurements of geometrical features of the maxillary sinus. The volume flux coming out of the sinus is

**Table 1:** Typical values of mucus properties and flow speeds 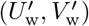 (top) and important dimensionless numbers (bottom), corresponding to mucociliary clearance in humans [28–30, 36–41]. Note that the value of 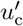 used to non-dimensionalise 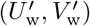 (and other quantities) in the main text corresponds to *µ* = 10^−1^ kg m^−1^ s^−1^ and *σ* = 0.08 N m^−1^ [30].

| Parameter | Description | Typical value | Units |
| --- | --- | --- | --- |
| $\rho$ | mucus density | $10^3$ | $\text{kg m}^{-3}$ |
| $\mu$ | mucus viscosity | $10^{-3}$ to $10$ | $\text{kg m}^{-1} \text{s}^{-1}$ |
| $\sigma$ | mucus-air surface tension | $0.01$ to $0.1$ | $\text{N/m}$ |
| $h'$ | mucus film thickness | $10$ to $15$ | $\mu\text{m}$ |
| $\ell'_s$ | sinus length-scale | $10^{-2}$ to $3 \times 10^{-2}$ | $\text{m}$ |
| $\epsilon = \frac{h'}{\ell'_s}$ | ratio of mucus film thickness to sinus length | $10^{-3}$ to $10^{-2}$ | dimensionless |
| $g$ | gravitational force per unit mass | $9.8$ | $\text{m s}^{-2}$ |
| $u'_c = \frac{\epsilon^2 \rho g \ell'^2_s}{\mu}$ | reference velocity scale | $10^{-3}$ to $10^4$ | $\mu\text{m s}^{-1}$ |
| $U'_w$ | tangential velocity at the wall | $1$ to $200$ | $\mu\text{m s}^{-1}$ |
| $V'_w$ | normal velocity at the wall | $10^{-5}$ to $10^{-2}$ | $\mu\text{m s}^{-1}$ |
| $\text{Bo} = \frac{\rho g \ell'^2_s}{\sigma}$ | Bond number | $\sim 10$ | dimensionless |
| $U_w = \frac{U'_w}{u'_c}$ | normalised tangential velocity | $10^{-1}$ to $40$ | dimensionless |
| $V_w = \frac{V'_w}{\epsilon u'_c}$ | normalised wall-normal velocity | $10^{-3}$ to $1$ | dimensionless |

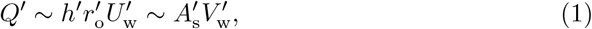

where *h*^′^ is the height of the mucus film, 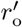 is the radius of the ostium and 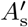 is the surface area of the maxillary sinus. The scaling in the first part of eqn. (1) follows from the assumption that in a healthy state, the mucus does not flow out through the total available ostium area (which would be proportional to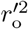), but only coats the inner surface of the ostium, forming a layer of thickness ∼ *h*^′^. The scaling in the second part of eqn. (1) follows from a mass balance argument that all the mucus secreted from the surface of the sinus must leave through the ostium. Now, if the volume of the sinus is 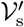 and its typical length-scale is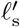, then its internal surface area is,

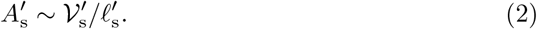

Combining eqns. (1) and (2) yields,

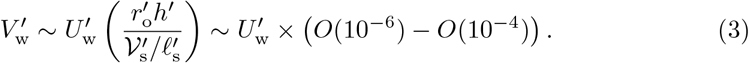

To arrive at the number in brackets in eqn. (3), we have used the following values of the geometric parameters, obtained from measurements on human sinuses: *h*^′^ ∼ 1015 *µ*m [14], 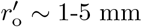 [13, 17, 31], 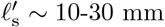 [17] and 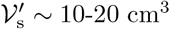[32, 33]. An estimate of 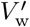 can also be made by dividing the volumetric rate of mucus production in the nasal epithelium, by the area of the nasal epithelium. Ref. [34] states that 20-40 mL of mucus is produced per day from around 160 cm^2^ of nasal mucosa; this yields an in-flow speed of 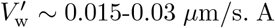. A third way to estimate 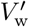 is by noting that ciliary beating causes turnover of the mucus blanket every 20–30 minutes [35]; so, if the thickness of the mucus film is 10-15 *µ*m, then 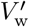 should be ∼ 5 × 10^−3^ *µ*m/s. In our theoretical study, we will cover a broad range of values of 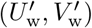 to reflect the wide variance in MCC rates across different sinus geometries and physiological conditions. We note that not all pairs of values of 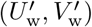 would correspond to the typical conditions inside a healthy sinus. The lowermost values of 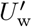 would reflect MCC in sinuses characterised by extensive cilia loss, whereas the largest values of 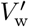 would be more representative of sinuses with mucosal swelling, a condition which leads to more mucus secretion [17].

## 3 Active, fluid-producing thin-film equations

The objective of our paper is to identify the physical conditions amenable to maintenance of a steady mucus layer inside the model sinus. We thus need to solve the equations governing mucus flow inside the sinus, and from them, deduce the shape of the mucus film. Since the typical thickness of the mucus layer *h*^′^ ∼ 10-15 *µ*m [14] is much smaller than the typical length-scale of the sinus 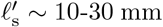 [17], the dynamics of mucus flow are governed by classical thin-film (lubrication) equations [42]. In this paradigm, the fluid’s velocity normal to the sinus walls is at least 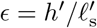 times smaller than its velocity along the sinus walls, where *ϵ* ≪ 1. Thus, the fluid flow is predominantly tangential to the sinus walls. In addition, the relative thinness of the mucus layer means that the variation of fluid velocity along the film is negligible as compared to its variation across the film. Finally, in the thin-film limit, the fluid pressure varies only along the film, while staying approximately constant normal to the film. These ideas are mathematically formalized in Appendices A.1 and B.1.

Under the simplifying assumptions listed above, a classical method may be used to derive the evolution equation satisfied by the mucus thickness [42]. One starts by expressing the (tangential) velocity of the fluid as a superposition of a pressure-driven flow resulting from variations in the height of the mucus film, a boundary-driven flow caused by the cilia-induced tangential velocity imposed along the cavity walls and a flow driven due to gravity. This velocity can then be used to calculate the tangential flux (i.e. flow rate) of mucus, as a function of the local height of the mucus film. Thereafter, one can use a mass balance argument to relate the rate-of-change of the mucus film’s height to the tangential mucus flux and the mucus production rate. In this way, the thin-film analysis allow us to reduce the multiple, coupled, nonlinear partial differential equations and boundary conditions describing the fluid’s flow-field, into a single nonlinear, partial differential equation describing the time evolution of the height of the mucus film [42]. While the mathematical details of the derivation of these thin-film equations are shown in Appendices A.1 and B.1, we provide here the final, dimensionless equations governing the film thickness, in both circular and spherical geometries.

### 3.1 Circular geometry

#### 3.1.1 Governing equation

In a symmetric system as shown in Fig. 2(a), the (dimensionless) thickness of the mucus film obeys (see Appendix A.1 for details),

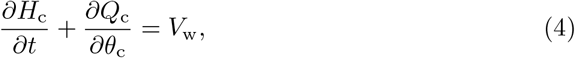

where,

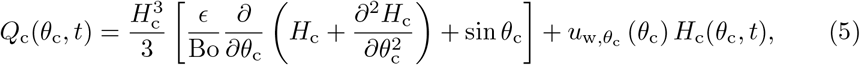

where the sub-script ‘c’ denotes circular geometry. In eqn. (4), *H*_c_(*θ*_c_, *t*) is the film thickness at location *θ*_c_ and time *t, Q*_c_ (*θ*_c_, *t*) is the local, tangential fluid flux and *V*_w_ is the dimensionless normal component of the fluid velocity at the cavity wall. In eqn. (5), 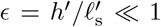 is the ratio of the characteristic film thickness *h*^′^ to the characteristic length-scale of the sinus 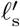; Bo is the Bond number, a dimensionless measure of the importance of gravity as compared to surface tension, in driving the film (see Appendix A.1). Also, *u*_w,*θ*c_ (*θ*_c_) in eqn. (5) is a prescribed tangential velocity at the walls of the circle (blue arrows in Fig. 2(a)), which models the action of the ciliated epithelium on the mucus; we discuss its functional form in Sections 3.1.3 and 3.3.

#### 3.1.2 Boundary conditions

The symmetry of the setup in Fig. 2(a) means that we just need to solve eqn. (4) over half the domain, i.e. for 0 ≤ *θ*_c_ ≤ *π*; where *θ*_c_ = 0 is the topmost point (orange dot in Fig. 2(a)) and *θ*_c_ = *π* is the bottommost point (black dot in Fig. 2(a)); the solution for *π* ≤ *θ*_c_ ≤ 2*π* can then be obtained by reflecting the solution for 0 ≤ *θ*_c_ ≤ *π* about the (vertical) axis. Symmetry also dictates that the flow-rate must vanish at *θ*_c_ = *π*, which yields the following conditions on *u*_w,*θ*c_ and *H*_c_:

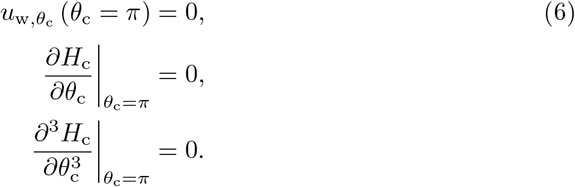

The first condition in eqn. (6) needs to be satisfied by design, by choosing a function *u*_w,*θ*c_ (*θ*_c_) that it is odd with respect to *θ*_c_ = *π* (see eqn. (10), Section 3.3). The second and third conditions follow from symmetry and the condition of continuity of film shape at *θ*_c_ = *π*.

#### 3.1.3 Modeling the ostium

For the circular geometry, we model the ostium as a discontinuity in the fluid velocity at *θ*_c_ = 0, or equivalently, at *θ*_c_ = 2*π*. Our choice of a symmetric ciliary wall-slip that is odd with respect to *θ*_c_ = *π* (shown qualitatively in Fig. 2(a); see also Section 3.3), disrupts the periodicity of a circular geometry at *θ*_c_ = 2*π*; in fact, it causes a jump, such that *Q*_c_ (*θ*_c_ = 2*π*) = −*Q*_c_ (*θ*_c_ = 0). However this is not a problem if we treat the point *θ*_c_ = 0, 2*π* as a local sink of fluid flow. Thus, once the action of the wall-slip causes the fluid to reach *θ*_c_ = 0, 2*π*, the fluid is instantaneously removed from the domain/cavity, much like mucus exiting the sinus from its ostium.

### 3.2 Spherical geometry

#### 3.2.1 Governing equation

For a spherical (but axisymmetric) geometry, the mucus film thickness satisfies (see Appendix B.1 for details),

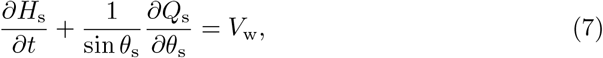

where,

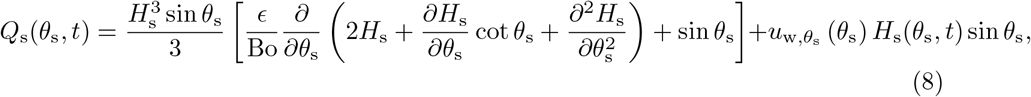

where the sub-script ‘s’ denotes spherical geometry. In eqn. (7), *Q*_s_(*θ*_s_, *t*) denotes the instantaneous, azimuthally averaged tangential flux at the location *θ*_s_ (i.e. the tangential flux normal to the dotted line in Fig. 2(b), averaged over the coordinate *ϕ*_s_). Similar to eqn. (5), *u*_w,*θ*s_ (*θ*_s_) in eqn. (8) is a prescribed tangential velocity at the walls of the spherical cavity.

#### 3.2.2 Boundary conditions

In the 3D axisymmetric case, symmetry dictates that we must have at the bottom pole (at *θ*_s_ = *π*),

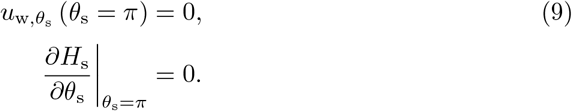

Once again the first condition in eqn. (9) needs to be satisfied by a suitable choice of *u*_w,*θ*s_ (see eqn. (10), Section 3.3). We do not need any other boundary conditions because the flux *Q*_s_ vanishes identically, by definition, at the bottom pole [26, 43].

#### 3.2.3 Modeling the ostium

In the 3D geometry, the fluid inside the cavity exits through an ostium modelled as a small, flat hole at the top, as marked by the orange circle in Fig. 2(b). Note that it is essential to truncate the sphere, and we cannot have a discontinuity-based exit from the top pole of an un-truncated/complete sphere; since for *θ*_s_ = 0 the flow-rate *Q*_s_ vanishes identically and so it is (understandably) impossible to exit as the radius of the orange ring in Fig. 2(b) tends to zero. In the present work, the radius of the model ostium is defined by an exit angle 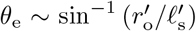 (see Fig. 2(b)), where 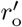 and 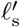 are, respectively, the typical ostium radius and the typical sinus length-scale. Using the values of 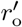 and 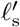 as mentioned in Section 2.3, we obtain *θ*_e_ ≈ 5° − 20°.

### 3.3 Physical description of the thin-film equations

Physically, eqns. (4) (2D) and (7) (3D) describe a mass-balance argument: the timerate-of-change of film-height at any section (*θ, t*), is the sum of the net fluid flux entering the section tangentially (−*∂Q*_c_*/∂θ*_c_ in eqn. (4) and − (sin *θ*_s_)^−1^ *∂Q*_s_*/∂θ*_s_ in eqn. (7)), and the fluid entering the section normally through the boundary, *V*_w_. Then, eqns. (5) (2D) and (8) (3D) describe the three contributions to the tangential fluid flux *Q*. The first is flow due to gravity, which is the term inside the square brackets that is proportional to sin *θ* (*θ* = *θ*_c_ or *θ*_s_) in eqns. (5) and (8). The second is the flow due to the effective action of the cilia, which is the last term in eqns. (5) and (8). The third contribution is the flow due to a surface-tension-driven pressure gradient resulting from spatial changes in the film’s curvature; this is the term multiplying *ϵ/*Bo in the square brackets.

The results in eqns. (4) and (5) in 2D (and, eqns. (7) and (8) in 3D) are extensions to the classical systems of equations governing thin-film dynamics over curved substrates [24–27], with two important additions: a wall-normal fluid velocity contribution *V*_w_ in eqns. (4) and (7), and an active tangential slip contribution *u*_w,*θ*c*/*s_ *θ*_c*/*s_ in eqns. (5) and (8). In our model, these represent respectively, the production of mucus inside the sinus, and the sweeping of the mucus toward the ostium by the ciliated cells. In the limits of *u*_w,*θ*c*/*s_ ≡ 0 and *V*_w_ ≡ 0, our formulation reduces to the classical (passive) formulations for cylinders [27] and spheres [26].

As mentioned above, we assume that the mucus enters the system at a uniform rate *V*_w_, normally at the wall. The spatial distribution of the tangential velocity, *u*_w,*θ*_ (*θ*), is motivated by the observation that “*mucociliary transport begins in the maxillary sinus as a star, from the bottom of the sinus and moves in various directions towards the ostium*” [44] (see also Fig. 1c). This is modelled, for both eqn. (5) and eqn. (8), by a hyperbolic tangent function,

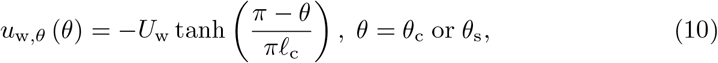

such that the tangential slip is zero at the bottom-most point (see the black dots at the bottom in Fig. 2) and increases to *U*_w_ over a relevant length-scale *ℓ*_c_, as we move up along the cavity. In the present work, we set *ℓ*_c_ = 0.5, for a smooth transition from 0 at the floor of the cavity, to ∼ *U*_w_ near the ostium; lower values of *ℓ*_c_, quantifying a more rapid spatial transition, have only a minor, quantitative effect on our main results. We note that for a circular geometry, this definition of *u*_w,*θ*c_ (*θ*_c_) leads to a discontinuity at *θ*_c_ = 0, 2*π*, such that *u*_w,*θ*c_ (*θ*_c_ = 0) = −*u*_w,*θ*c_ (*θ*_c_ = 2*π*), but, as explained in Section 3.1.3, this is not a problem because *θ*_c_ = 0, 2*π* denotes a fluid sink for the 2D geometry, and hence allows for discontinuity of the wall velocity.

### 3.4 Numerical solution and validation

We numerically solve eqns. (4) and (7), with the boundary conditions (6) and (9) respectively, using a semi-implicit finite-difference method whose details are provided in Appendices A.2 and B.2. We validate our numerical solution in the limit of zero mucus production (*V*_w_ ≡ 0) and sweeping (*U*_w_ ≡ 0), by reproducing classical results of the drainage of a thin film over a cylindrical [27] and a spherical [26] substrate, as shown in Figs. A2a and B3a in Appendices A and B, respectively.

## 4 Steady mucus drainage in active fluid-producing thin films

We begin our results with a comment on the dimensionless values of 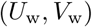, whose corresponding dimensional values 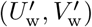 were discussed in Section 2.3. A natural velocity scale in the present problem is set by gravity, 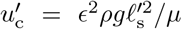(see Appendix A.1), where *µ* is the fluid’s dynamic viscosity, whose range of values is given in Table 1. This is the characteristic velocity with which a thin film would flow down a substrate due to gravity alone. In the thin-film analysis, the fluid velocities tangential and normal to the surface are made dimensionless using 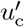 and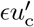, respectively, which yields (with *ϵ* = 10^−3^ and 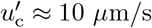; see Table 1):

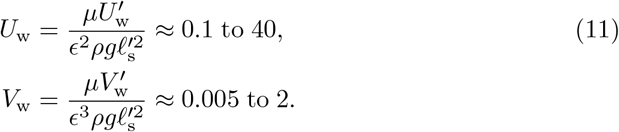

### 4.1 Mucus film evolution in two dimensions

We illustrate in Fig. 3 two representative examples of the time evolution of the (thin) mucus film in a circular cavity, i.e. in two dimensions. In Figs. 3a (Cartesian plot) and 3b (polar plot), the active wall-slip (ciliary action) is not sufficiently strong to push out the fluid that is being produced in the cavity walls. Hence, the fluid inside the sinus increases in volume with time and, due to gravity, it accumulates at the bottom. This results in a progressive increase in the film height at the bottom of the cavity, until the thin-film approximation breaks down and the situation becomes non-representative of mucus flow inside sinuses. However, if the magnitude of the tangential slip, *U*_w_, is increased beyond a threshold, then one does obtain a steady solution, as shown in Fig. 3c and 3d. In this case, the active motion (*U*_w_) is sufficiently large to overcome gravity; it then drives the fluid out of the cavity and balances the local fluid production (*V*_w_), leading to the development of a thin mucus layer, as is expected inside healthy sinuses.

**Fig. 3:**
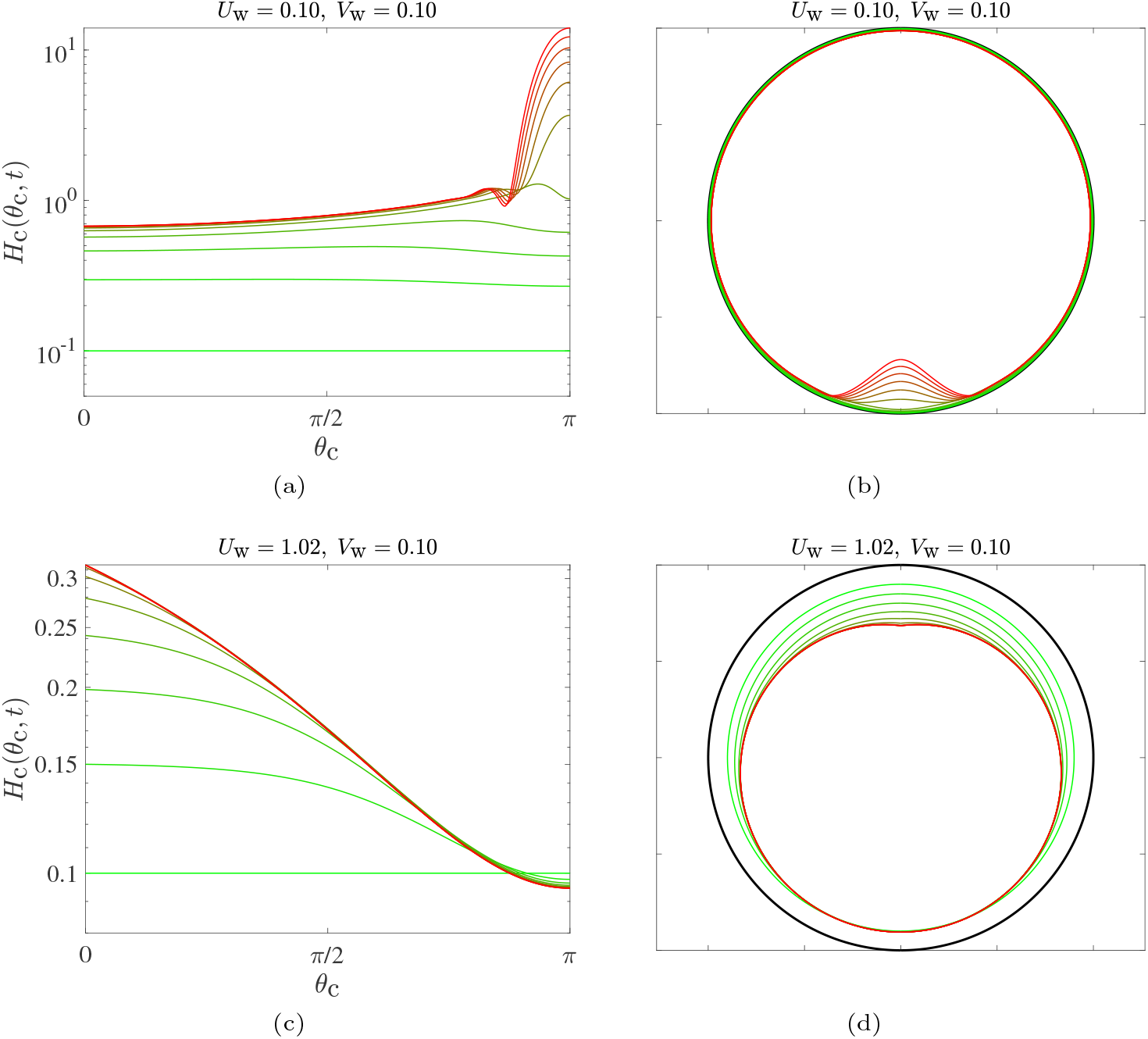
The two regimes for the time evolution of the thin (mucus) film inside a circular cavity. (a-b) Time evolution of the film for *U*_w_ = 0.10 and *V*_w_ = 0.10, for which eqn. (4) does not have a steady solution; panel (a) is a Cartesian plot, and panel (b) is a polar plot where the film thickness has been magnified 20 times the actual value, to help visualisation. The profiles evolve from dimensionless time *t* = 0 (green) to *t* = 20 (red) in time intervals *Δt* = 2. (c-d) Time evolution of the film inside the circular cavity for *U*_w_ = 1.02 and *V*_w_ = 0.10, for which eqn. (4) reaches a steady solution; panel (c) is a Cartesian plot and panel (d) is a polar plot where the film thickness has been magnified 1000 times the actual value. The profiles evolve from dimensionless time *t* = 0 (green) to *t* = 5 (red) in time intervals *Δt* = 0.5. For these set of results, we considered *ℓ*_c_ = 0.3.

For the cases where a steady thin film can be obtained (i.e. when *U*_w_ is sufficiently large), the shape of the film as a function of the wall-slip, is shown in Fig. 4a. As expected from intuition, larger values of the characteristic slip *U*_w_, result in thinner films (for a fixed rate of fluid injection *V*_w_). We can obtain an expression for the exit-height of the film, *H*_c_(0), in terms of (*U*_w_, *V*_w_) by integrating eqn. (4), ignoring the contribution from the *ϵ/*Bo term (since *ϵ* ≪ 1), and noting that in steady state,

**Fig. 4:**
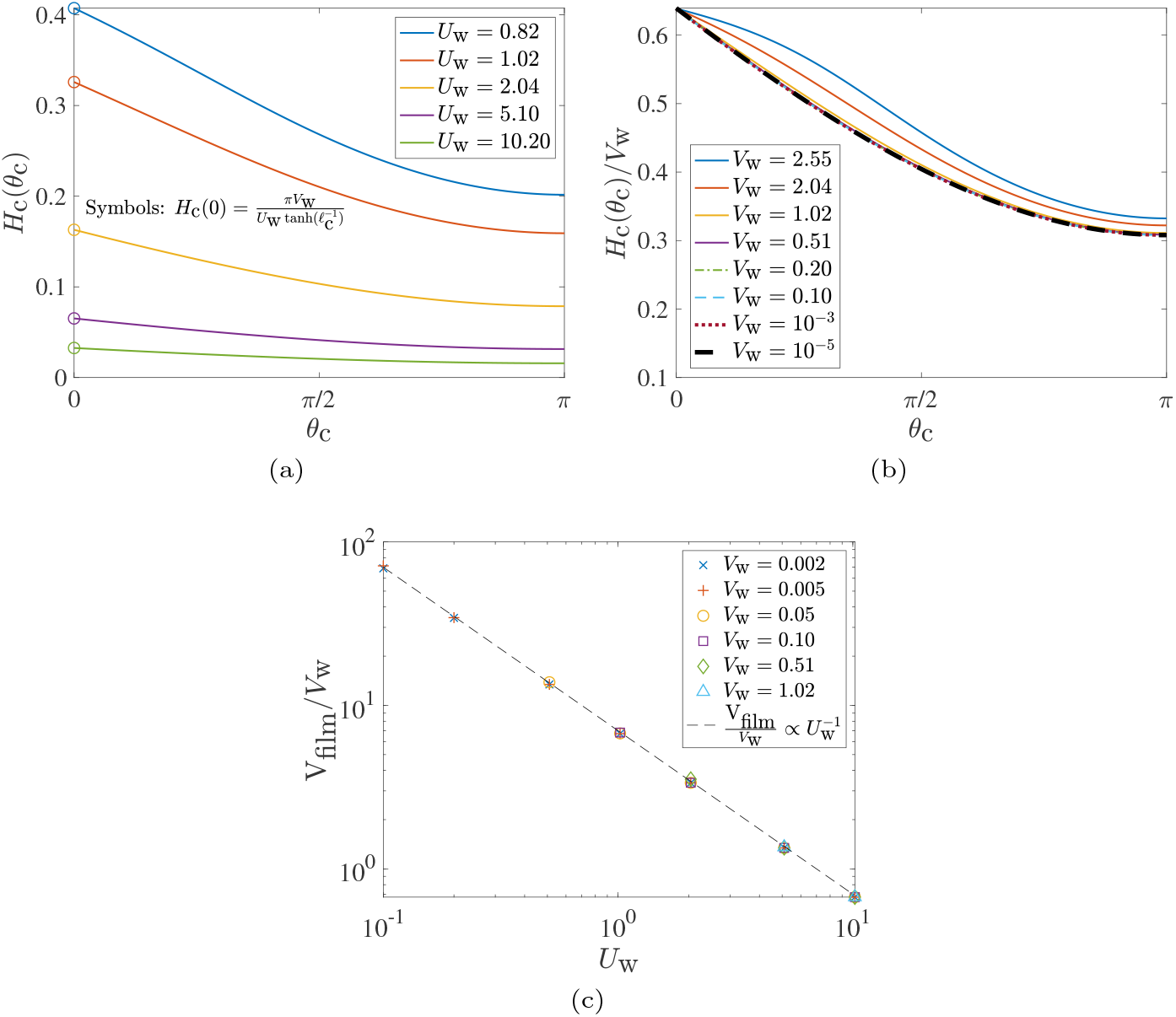
Height of steady-state film in the two-dimensional geometry as function of active parameters *U*_w_ and *V*_w_. (a) Variation of the film height for a twodimensional/circular cavity, as a function of the effective ciliary clearance speed, *U*_w_, for a fixed mucus injection rate *V*_w_ = 0.10. (b) Steady-state film height normalised by the rate of mucus injection, *H*_c_(*θ*_c_)*/V*_w_, for different values of the injection rate; *U*_w_ = 5.10 for all the plots. (c) Scaling of the film volume, *V*_film_, with the injection rate, *V*_w_, and the speed of ciliary clearance *U*_w_, for low values of the injection rate.

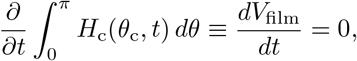

where, *V*_film_ is the volume of the mucus film. This yields,

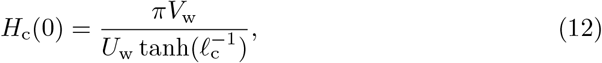

a prediction that is indeed confirmed by our numerics, as shown by the circles in Fig. 4a. Interestingly, eqn. (12) tells us that the steady-state exit-height in our problem does not depend on the fluid’s properties (via the Bond number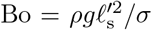) and depends only on the specified kinematics through (*U*_w_, *V*_w_, *ℓ*_c_). Since it was necessary to neglect the *ϵ/*Bo term in order to arrive at eqn. (12), this means that surface tension plays a negligible role in film dynamics for the cases where a steady solution exists to eqn. (4).

In Fig. 4b we next show the normalised steady-state film shape, *H*_c_(*θ*_c_)*/V*_w_; of course, such a representation is valid only for *V*_w_ ≠ 0. It is clear that the average film-thickness increases monotonically with increasing *V*_w_. For the lower-most values of *V*_w_ considered, the steady-state plots of *H*_c_(*θ*_c_)*/V*_w_ collapse onto each other; this is true for *V*_w_ as low as 10^−5^. One may then write, *H*_c_(*θ*_c_) ≈ *V*_w_ × *f* (*θ*_c_; *U*_w_, *ℓ*_c_), for a large range of mucus production rates: 10^−5^ ≤ *V*_w_ *≲ O*(1).

If we postulate that the film volume *V*_film_ is proportional to the exit height *H*_c_(0), then based on eqn. (12) we may conclude that the normalised film volume, *V*_film_*/V*_w_, is inversely proportional to the ciliary slip *U*_w_; this is indeed confirmed numerically in Fig. 4c. We thus obtain a scaling estimate of the amount of mucus maintained inside the two-dimensional cavity, for rates of mucus injection that admit a steady solution over a large range of effective ciliary clearance strengths.

### 4.2 Mucus film evolution inside a sphere (three dimensions)

We now consider the three-dimensional case and show in Fig. 5 the time evolution of the mucus film inside the spherical cavity. The parameters in Figs. 5a and 5b correspond to the case where *U*_w_ is not sufficiently large to overcome gravity and Figs. 5c and 5d corresponding to the case where the active flow *U*_w_ is strong enough that a steady state can be reached. Both the unsteady and steady-state film shapes in the spherical case are qualitatively different from the circular case and there is a sharper increase in the film height (toward the bottom for the unsteady solutions in Figs. 5a and 5b, and also toward the top for the steady solution in Figs. 5c and 5d). In particular, the steady-state mucus film collects fluid as it develops from the bottom to the top of the cavity; and since the fluid must exit from a narrow constriction at the top, the film thickens much more rapidly than in the circular case.

**Fig. 5:**
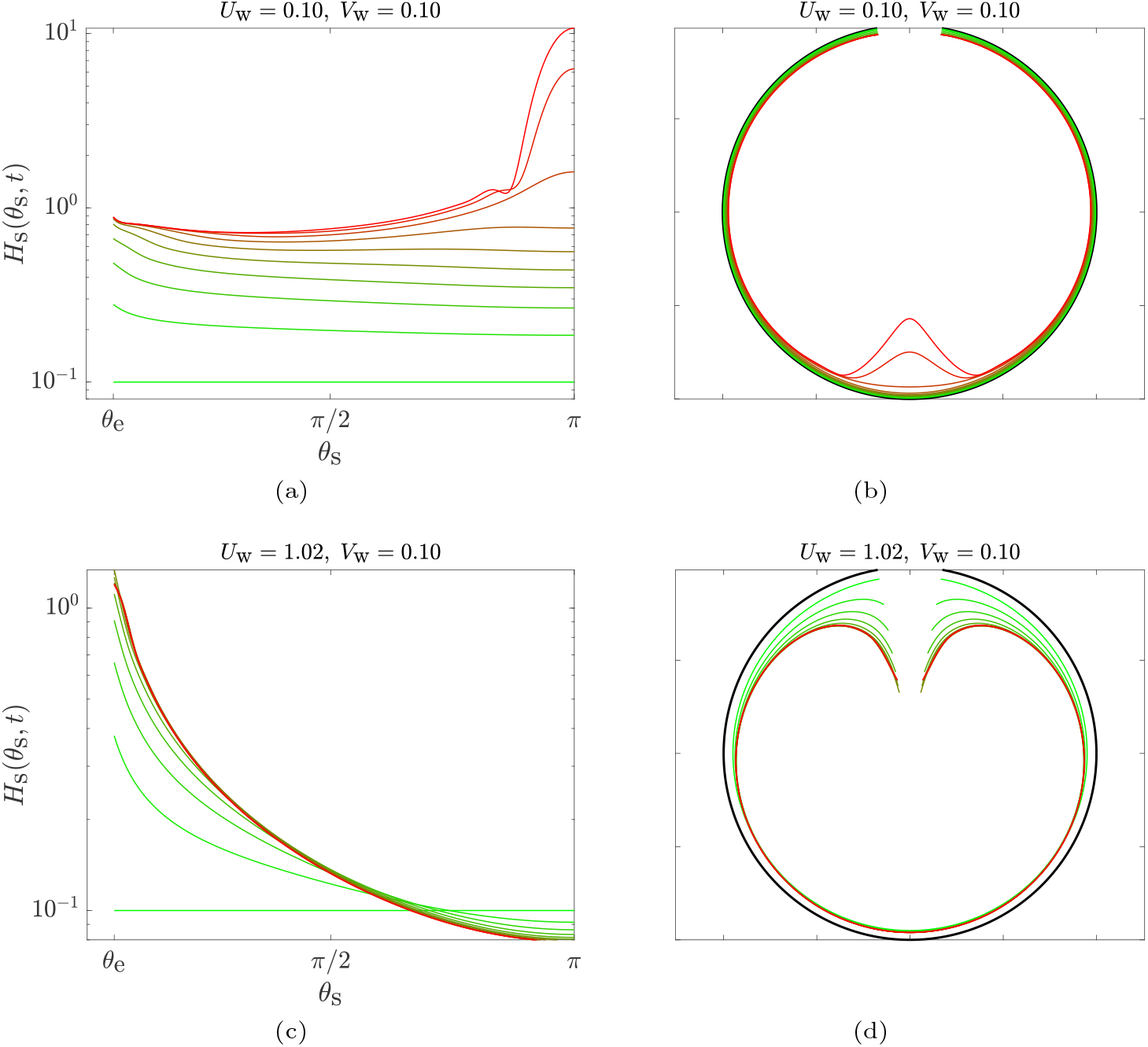
Time evolution of the film inside the spherical cavity. (a-b) Case with *U*_w_ = 0.10 and *V*_w_ = 0.10, for which eqn. (7) does not have a steady solution; panel (a) is a Cartesian plot, and panel (b) is a polar plot where the film thickness has been magnified 40 times the actual value, for visualisation purposes. The profiles evolve from dimensionless time *t* = 0 (green) to *t* = 9 (red) in time intervals *Δt* = 1. (c-d) Case with *U*_w_ = 1.02 and *V*_w_ = 0.10, for which eqn. (7) reaches a steady solution; panel (c) is a Cartesian plot, and panel (d) is a polar plot where the film thickness has been magnified 500 times the actual value. The profiles evolve from *t* = 0 (green) to *t* = 4.80 (red) in time intervals *Δt* = 0.4. For these set of results, we considered *ℓ*_c_ = 0.5.

The steady-state exit height, denoted by *H*_s_(*θ*_e_), is related nonlinearly to (*U*_w_, *V*_w_) via,

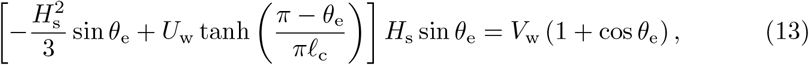

which can be derived by ignoring the surface tension contribution in eqn. (7) (because *ϵ/*Bo ≪ 1), multiplying its steady version by sin *θ*_s_ and integrating from *θ*_s_ = *θ*_e_ to *θ*_s_ = *π*. For *H*_s_(*θ*_e_) *≲ O*(1) and *θ*_e_ ≪ 1, eqn. (13) yields,

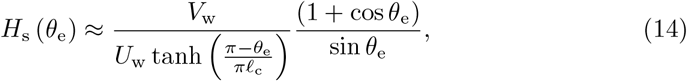

which is compared against the numerical results in Fig. 6, where we see that the analytical prediction best matches the numerical results for the thinner films and a mismatch occurs mainly when the exit height is not small, *H*_s_(*θ*_e_) ∼ *O*(1). For (*U*_w_ = 5.10, *V*_w_ = 1.02) in Fig. 6b, eqn. (14) overestimates the exit height because it ignores the contribution from surface-tension-induced pressure gradients. The latter become important near the exit, where rapid mucus accumulation results in sufficiently large gradients in the mucus film thickness, causing surface-tension-driven flows that reduce the exit height. Note that this role of surface tension is unique to the spherical geometry and is not seen for the circular geometry. The analytical estimate of the exit height for the circular geometry (eqn. (12)) also ignored surface tension, but it matched perfectly with the numerical results for a wide range of (*U*_w_, *V*_w_) (Fig. 4a). Thus, for the biologically-relevant values listed in Table 1, surface tension effects are truly negligible for the 2D/circular geometry, but this is not always the case for the 3D/spherical geometry.

**Fig. 6:**
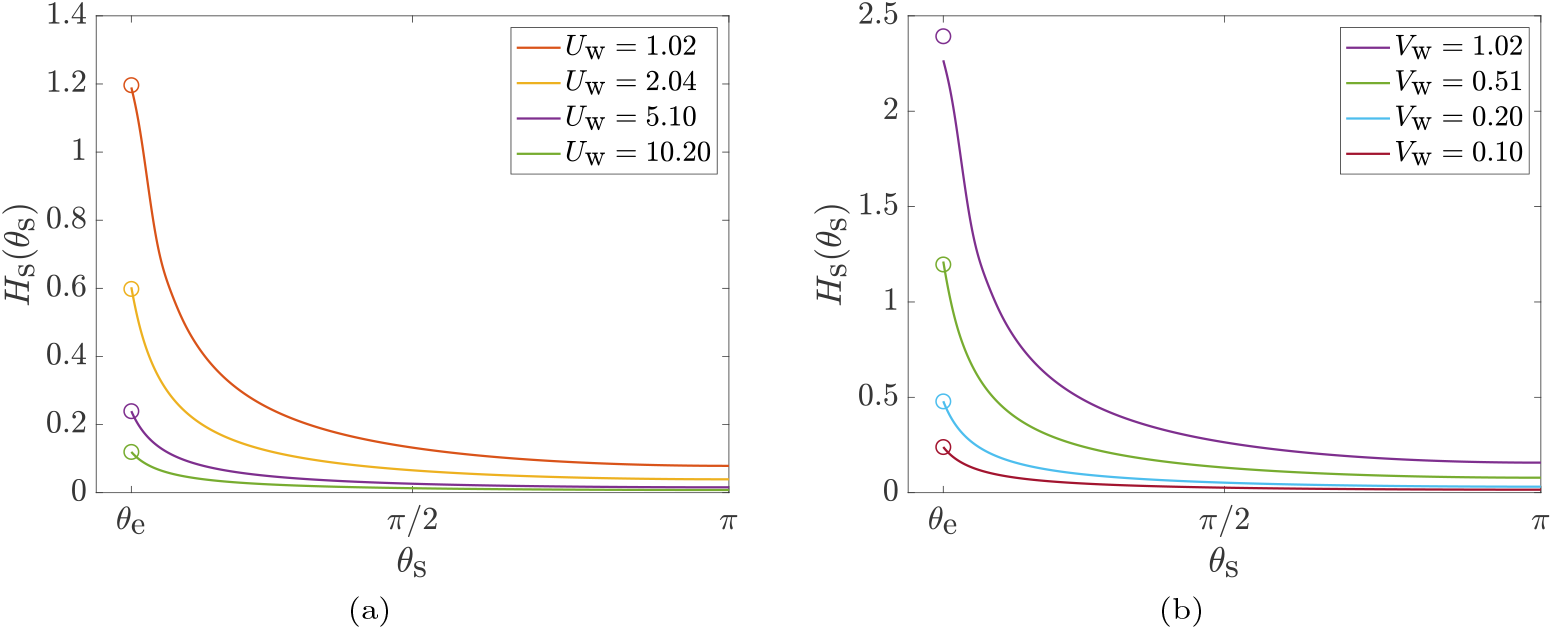
Height of steady-state film in the three-dimensional geometry as function of active parameters *U*_w_ and *V*_w_. (a) Film height as a function of the effective ciliary clearance speed, *U*_w_, for a fixed mucus injection rate *V*_w_ = 0.10. (b) Film height as a function of the mucus injection rate, *V*_w_, for a fixed effective ciliary clearance speed *U*_w_ = 5.10. The circles denote the analytical estimate of the exit height, based on eqn. (14).

## 5 Existence of a steady solution

### 5.1 Phase space of solutions

In the previous sub-section, we demonstrated that depending on the relative values of (*U*_w_, *V*_w_), the mucus film either builds up at the bottom of the cavity, or attains a steady-state shape wherein the mucus is cleared from the cavity at the same rate that it is produced at the cavity walls. This was the case both in two and three dimensions. Using our numerical model, we can systematically vary the two active parameters, *U*_w_ and *V*_w_, and map out the existence of these two different solutions. The results are shown in Fig. 7a for a circular (2D) cavity and in Fig. 7b for a spherical (3D) geometry with exit angle *θ*_e_ = 5°.

**Fig. 7:**
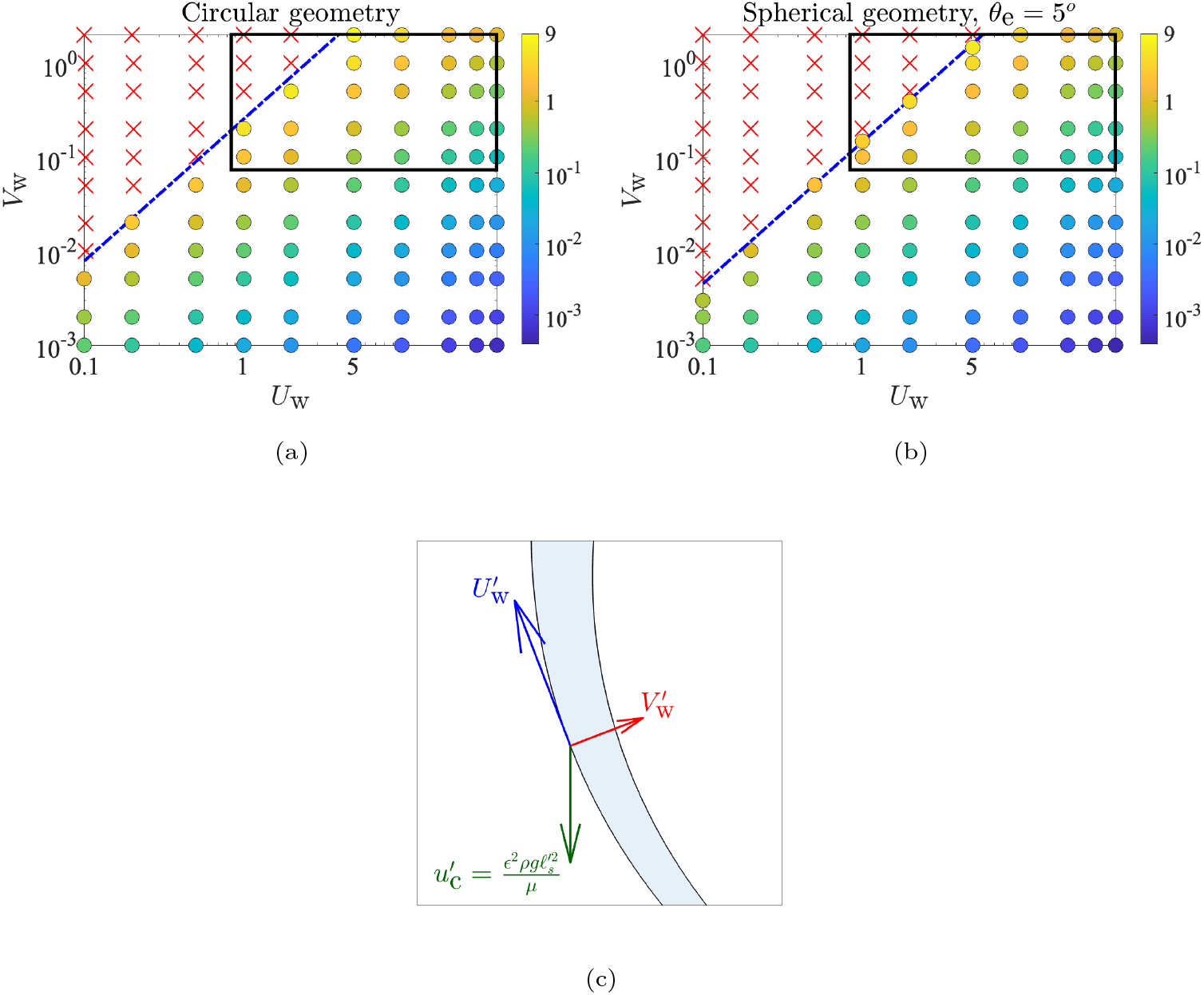
Phase space of steady (·) vs unsteady (×) solutions as a function of (*U*_w_, *V*_w_) for, (a) the circular geometry and (b) spherical geometry with *θ*_e_ = 5°. The red crosses (×) denote cases where mucus accumulates inside the cavity, whereas the coloured circles (·) denote cases where a steady mucus layer is formed, with colours quantifying the steady-state film volume normalised by the initial film volume. The blue line represents the transition scaling 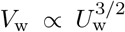 as predicted by eqn. (18). The black rectangle denotes the estimated range of values of *U*_w_ and *V*_w_ for human sinuses in healthy conditions. (c) Sketch of a magnified view of the mucus film and the three relevant velocities that govern the evolution of its shape.

As expected, a steady solution exists whenever the rate of mucus in-flow (*V*_w_) is particularly low, or the effective ciliary velocity (*U*_w_) is sufficiently high. The principal effect of the cavity geometry (circular versus spherical) is reflected in the slightly larger region of existence of steady solutions for the circular case. However, the general shape of the boundary demarcating steady and unsteady solutions remains unchanged between the circular and the spherical case. This suggests that the existence of a steady solution is due to the same fundamental physics in both geometries, which we rationalise below.

### 5.2 Steady vs unsteady solutions: Scaling analysis

We now estimate the relation between *U*_w_ and *V*_w_ which defines the boundary between the steady and unsteady solutions in Figs. 7a and 7b, i.e. we derive a scaling between *U*_w_ and *V*_w_ for which eqns. (4) and (7) are expected to admit a steady solution.

We start by a sketch of a typical section of the film, shown in Fig. 7c, highlighting the three relevant velocity scales governing the shape of the mucus film: fluid is produced at the walls at a rate 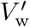, from where its motion is governed by a competition between a typical gravitational drainage velocity 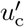 and an effective ciliary velocity 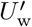, which tries to drive the fluid up and out of the cavity. Conservation of mass in the classical thin-film limit sets the relative scaling of 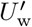 and 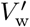 as,

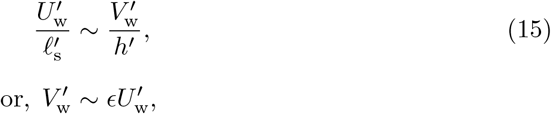

where we have used 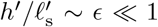. Further, we argue that the active (ciliary) wallvelocity 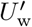 must be greater than the characteristic gravitational velocity scale 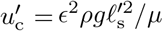, in order to successfully drive the mucus out of the cavity, meaning, we require

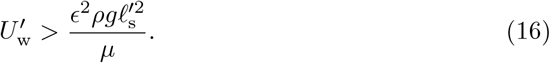

The scalings in Eqns. (15) and (16) can be combined to yield,

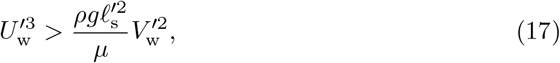

which can be non-dimensionalised using the appropriate velocity scales in the thin-film limit (see beginning of Section 4 and eqn. (11)) to obtain,

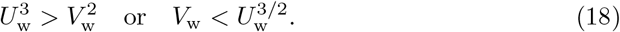

The resulting scaling 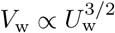 from eqn. (18) has been plotted in Figs. 7, where we see that it aligns well with the boundary demarcating the unsteady solutions from the steady solutions, for both the circular (Fig. 7a) and the spherical system (Fig. 7b). Thus, the threshold clearance velocity required to obtain a steady mucus layer, say 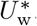, scales as the 2/3^rd^ power of the rate of mucus in-flow, i.e. 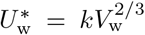 (by inverting eqn. (18)), where the constant *k* can be determined from numerical solutions to eqns. (4) and (7).

## 6 Application to mucociliary clearance in human sinuses

Using our theoretical model, we have identified the hydrodynamic conditions, specified by values of (*U*_w_, *V*_w_), under which a steady mucus layer can exist inside the cavity. Based on the discussions in Section 2.3, the operative conditions inside a healthy sinus correspond to an effective ciliary velocity, 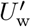 in the range 30 to 400 *µ*m/s and the mucus in-flow 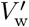 in the range 5 × 10^−3^ to 3 × 10^−2^ *µ*m/s. Using the characteristic velocity scales defined in the beginning of Section 4, the dimensionless values of the operative ciliary velocity and the mucus in-flow rate are thus given by *U*_w_ ∼ 1−40 and *V*_w_ ∼ 0.5−3. The region of the solution space that lies within this range is shown as rectangles in Figs. 7a and 7b. We see that, in general, these values do correspond to the existence of a steady solution according to our model. We thus postulate that the primary factors responsible for maintaining a steady mucus layer inside a healthy sinus are a combination of (i) the rate of mucus flow due to ciliary beating being sufficiently fast to overcome local gravitational drainage, and (ii) the rate of mucus production per unit area of the sinus being sufficiently small (as compared to the rate of ciliary clearance). Diseased conditions, such as excessive cilia loss or mucosal inflammation, violate one or both requirements, and thus, according to our model, will not lead to the formation of a thin mucus film over the sinus [17].

It is estimated that it takes 20–30 minutes to replenish the mucus film during MCC, although this time varies significantly, even in healthy individuals [35]. We may use our model to compute the time 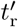 taken to reach the steady-state from an initially small film height, *H*_s_(*t* = 0) = 10^−3^, for values of (*U*_w_, *V*_w_) that fall within the physiological range outlined in Fig. 7b. This is illustrated for three cases in Fig. 8 and that time is seen to vary from 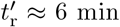 (when (*U*_w_ ≈ 40, *V*_w_ ≈ 2)) to 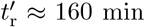 (when (*U*_w_ ≈ 1, *V*_w_ ≈ 0.1)). For (*U*_w_ ≈ 10, *V*_w_ ≈ 1), values that lie in the middle of the physiological range, we obtain 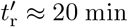. Thus, in addition to predicting the healthy operating conditions, our model is also able to approximately recover the typical mucus turnover rates observed in humans, under normal conditions.

**Fig. 8:**
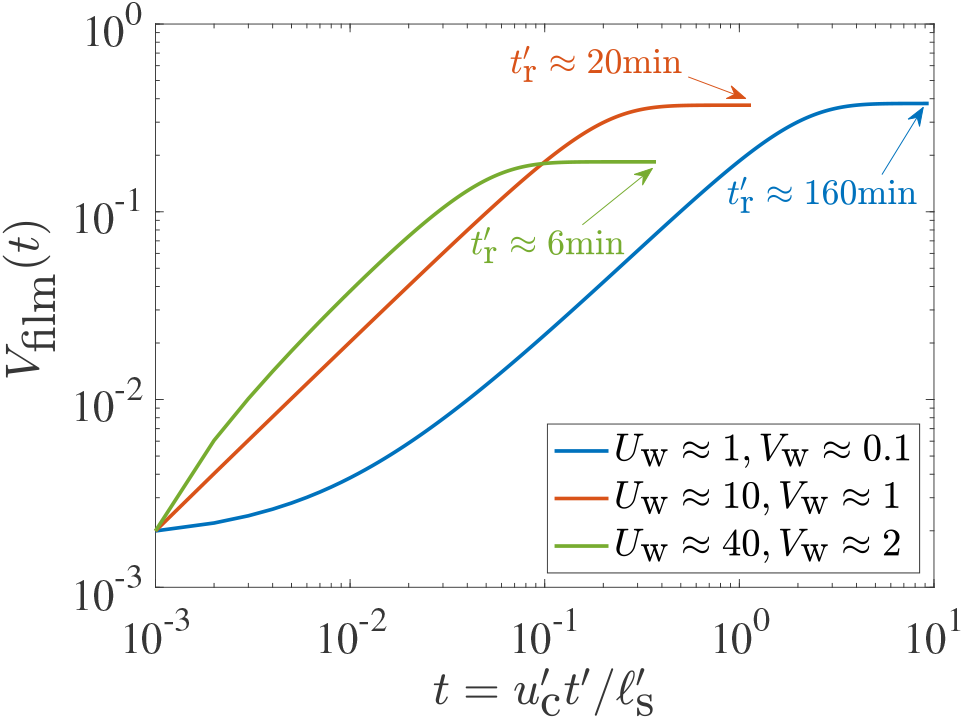
Time-evolution of film volume until steady state is reached, for three values of the pair (*U*_w_, *V*_w_). The steady-state is considered to have been reached when the absolute rate of change of film volume falls below a threshold,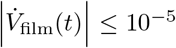. The dimensional time at which the steady-state is reached, denoted by 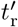, is indicated for each case. Note that the horizontal axis shows the dimensionless time; the characteristic time-scale is given by 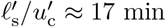.

An important geometric factor that affects mucus transport out of the sinuses is the size of the sinus opening, or the ostium. By varying *θ*_e_ in our 3D model, we can obtain further insight on the influence of the ostium size on mucus clearance. Typical ostium diameters range from 2-10 mm [13, 17, 31], which means that for a characteristic sinus length-scale 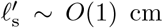 [17], the exit angle *θ*_e_ ranges from 5°-20°. The solution space for *θ*_e_ = 5° is compared with that for *θ*_e_ = 20° in Fig. 9. We see that there do exist instances in the (*U*_w_, *V*_w_) space where the fluid/mucus does not get cleared from the cavity with the narrower opening but it does get cleared from the cavity with a larger opening; these are identified in Fig. 9 by the filled red squares (representing results for *θ*_e_ = 5°) which coincide with the empty green circles (representing results for *θ*_e_ = 20°). Overall, however, an increase in the ostium radius is seen to cause only a modest change in the nature of the solution space.

**Fig. 9:**
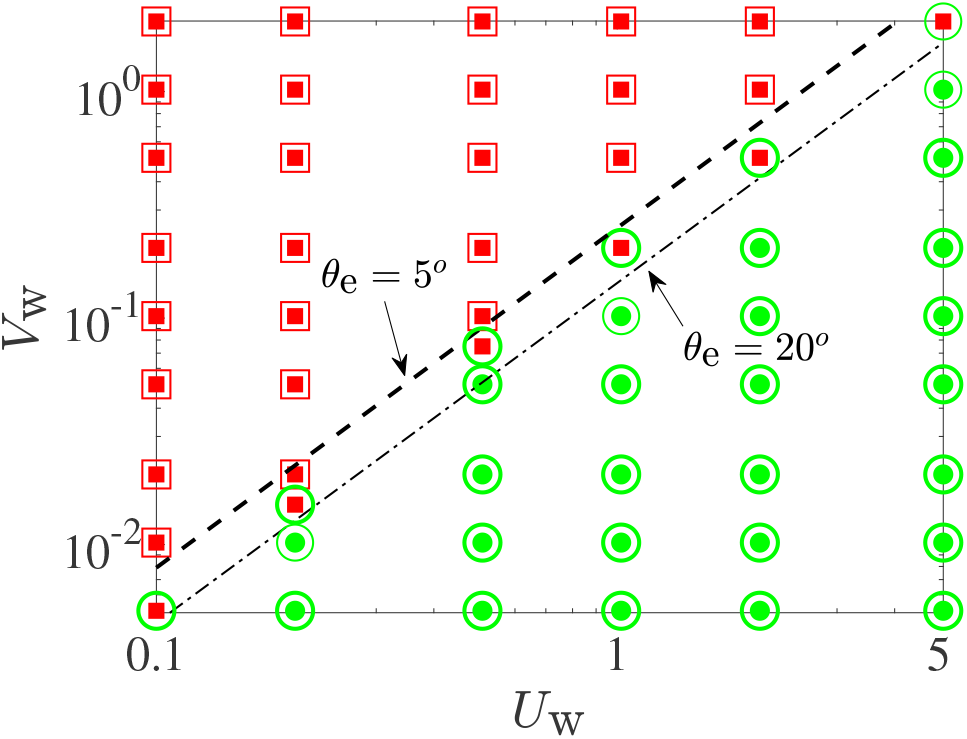
Effect of varying ostium size, quantified by the exit angle *θ*_e_ (see Fig. 2(b)), on the solution space in 3D model. Empty symbols are used to denote the solution type for the case with the broader exit angle (*θ*_e_ = 20°) whereas filled symbols denote the solution type for the case with the narrower exit angle (*θ*_e_ = 5°). There exists a small range (between the thin dash-dotted line and the thick dashed line) where solutions for *θ*_e_ = 5° (filled red squares) are unsteady but the solutions for *θ*_e_ = 20° (empty green circles) are steady.

Interestingly, diseased sinuses appear to be accompanied by other pathologies such as nasal polyps, which are benign, painless growths in and around the sinuses that obstruct mucociliary clearance by blocking the ostium. This condition can be treated by surgically removing the polyps, unblocking the ostium and restoring smooth mucus flow out of the sinuses. Our model also hints at the efficacy of polyp-removal surgeries: it shows that an increase in the size of the ostium from *θ*_e_ = 5° to *θ*_e_ = 20° (see Fig. 2(b)) doubles the maximum value of the mucus production rate, say *V*_w,max_, for which a steady mucus layer can exist inside the cavity. For example, Fig. 9 shows that, for *U*_w_ ≈ 2, *V*_w,max_ ≈ 0.2 when *θ*_e_ = 5° but it increases to *V*_w,max_ ≈ 0.5 when *θ*_e_ is increased to 20°. Similar 2-fold increments in *V*_w,max_ can be seen for other values of *U*_w_ as well, whenever *θ*_e_ is increased from 5° to 20°.

## 7 Conclusion and Perspectives

### 7.1 Summary of modelling

We considered in this paper the problem of thin-film fluid flow inside circular (2D) and spherical (3D) cavities, as a model for active mucociliary clearance (MCC) in the maxillary sinuses. Building on classical work for passive thin films, we derived a new nonlinear, partial differential equation for the time evolution of a thin film of fluid (mucus) that is released from the walls of a cavity (sinus) and driven, against gravity, toward an exit (ostium) by ciliary pumping, which is modelled as a prescribed tangential velocity at the cavity walls (active slip). Numerical solutions to this equation reveal two different behaviours in the long term: the mucus can either build up progressively at the bottom of the cavity or be cleared out at the same average rate with which it is produced, leading to the formation of a thin, steady film lining the cavity. These two regimes are demarcated on a phase-space of solutions (see Figs. 7a in 2D and 7b in 3D) defined by the rate of mucus production (denoted, in dimensionless form, as *V*_w_) and the rate of mucus clearance by cilia (*U*_w_, in dimensionless form). The fate of the mucus is decided by the relative magnitudes of *U*_w_ and *V*_w_. Using a scaling analysis based on physical arguments, we showed show that the threshold clearance velocity required to obtain a steady mucus layer scales as 2/3^rd^ power of the rate of mucus in-flow, i.e. the line separating the steady and unsteady solutions in Figs. 7a and 7b is given by 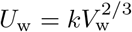, with a constant *k* that depends on the system geometry.

### 7.2 Summary of biological relevance

Biologically, mucus is produced in the sinuses at a rate 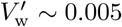 to 0.03 *µ*m/s, due to hydration of mucins secreted by goblet cells. The cilia push this mucus out of the sinuses with a velocity in the range 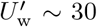 to 400 *µ*m/s. For typical values of the physical properties of mucus (see Table 1), the intrinsic gravitational drainage/settling velocity is 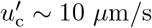. These values correspond to a healthy sinus, and hence they must lead to emergence of a steady state in our model system. This is indeed the case, most notably for the larger values of 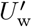, as shown in Fig. 7. Our theoretical model thus captures the essential physical ingredients responsible for successful mucociliary clearance, particularly in the maxillary sinus, where it is known that the cilia must work against gravity to deliver mucus to the nasal cavity [17, 21–23].

### 7.3 Model extensions

Our model uses many assumptions, which could be relaxed in future studies. Firstly, the ostium of the maxillary sinus isn’t always located at the highest point in the cavity and is often located on a medial wall [17]. In terms of the present model, this would amount to a rotation of the gravity vectors shown in Fig. 2, leading to loss of axisymmetry in the spherical case. When the ostium is not located symmetrically as shown in Fig. 2(b), one can develop and solve a non-axisymmetric thin-film equation for the time evolution of the film height as a function of the polar (*θ*_s_) and azimuthal (*ϕ*_s_) angles. This would require a conceptually straightforward, albeit numerically cumbersome, extension of the current work; where a key step would be to identify the form of the ciliary slip, *u*_w,*θ*s_ (*θ*_s_, *ϕ*_s_) (see eqn. 8).

Secondly, we treat the mucus as a single Newtonian fluid, whereas in reality it is a bi-layered, viscoelastic and shear-thinning fluid [9, 10]. The non-Newtonian rheology of the mucus will cause it to react differently to the ciliary slip than a Newtonian (purely viscous) fluid. These effects may significantly change the structure of the thin film equations (eqns. (4)-(8)), hence the shape of the mucus film inside the cavity and likely the phase-space of solutions in Fig. 7.

Thirdly, the maxillary sinus has a very complex geometry that isn’t fully captured by any one regular shape. It is often described to be pyramidal, and characterised by geometrical features such as recesses and protrusions [17]. Hence, an investigation of the influence of the actual sinus shape on MCC must extend the current work to cavities containing one or more of these features. Initial progress along this direction can be made for shapes that are small deviations from a sphere/circle, but analysis for more realistic shapes would necessitate the use of extensive computations.

The agreement between our predictions of steady-state operating conditions in sinuses and existing estimates of mucociliary clearance rates (Fig. 7b), shows that our model successfully captures the key physical mechanism responsible for uninterrupted mucus flow in the sinuses, and is thus encouraging. However, the simplicity of our model can restrict certain quantitative comparisons with real systems, for example, on aspects related to spatial variation of the film shape and the total volume of mucus contained in the film. Thus, further investigations of mucociliary clearance in sinuses is warranted to fully explore the appropriate physical conditions required to maintain healthy sinuses.

## Acknowledgements

We thank Bartlomiej Waclaw for useful comments. This work was funded by EPSRC (grant EP/W024012/1 to EL).

## Data availability

The results in this manuscript are based on theoretical derivations and not on existing experimental data. All steps of the derivations are given in the manuscript’s Appendix and thus can be used/reproduced as such.

## Appendix A 2D/circular system

**Fig. A1:**
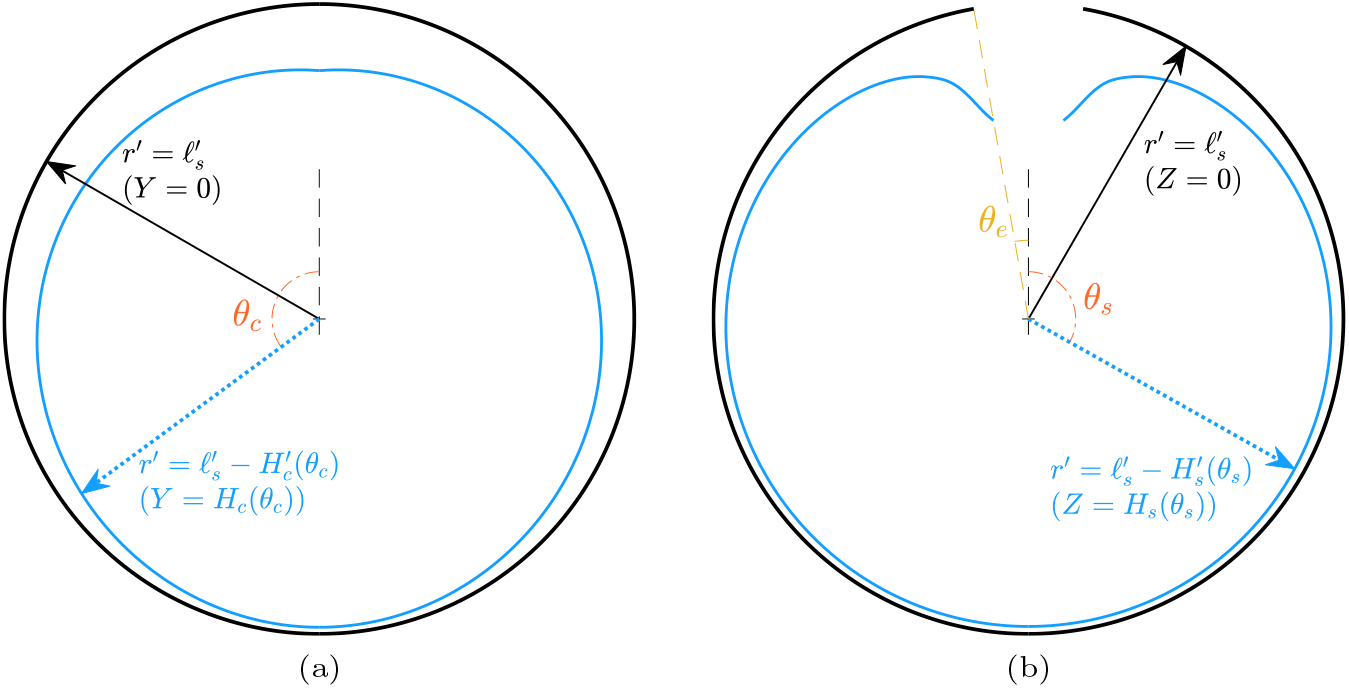
Sketch of the dimensional coordinate system used to describe (a) the circular cavity/sinus, and, (b) the spherical cavity/sinus. The black, solid arrow identifies the walls (black circle) and the blue, dotted arrow identifies the free surface (blue curve) of the fluid/mucus. Note that panel (a) is a planar/2D geometry (0 ≤ *θ*_c_ ≤ 2*π*), whereas panel (b) is a section of an otherwise 3D geometry (*θ*_e_ ≤ *θ*_s_ ≤ *π*); see also Fig. 2.

### A.1 Derivation of the thin-film equation

The fluid flow inside the sinus is dominated by viscous forces (i.e. the inertia of the fluid is negligible), and hence is governed by the Stokes equations and the incompressibility condition (i.e. continuity equation) [42]. For the 2D/circular system, we will work in polar coordinates (*r*^′^, *θ*_c_, *z*^′^). Since we are interested in a planar flow, the *z*^′^-component of the velocity and all derivatives with respect to the *z*^′^-coordinate are identically zero, i.e. 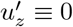 and also *∂*()*/∂z*^′^ ≡ 0.

The system geometry is described in Fig. A1a, where the walls of the cavity/sinus are at 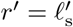 and the free surface of the fluid/mucus film is at 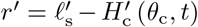. The starting point for deriving eqn. (4) is to non-dimensionalise the equations governing fluid flow in polar coordinates using the following reference scales:

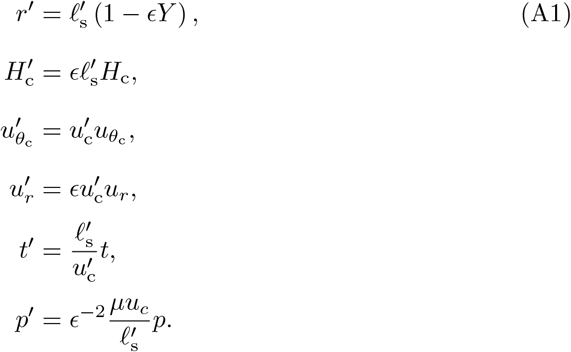

In the above, 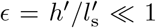 is a small parameter defined as the ratio of the typical thickness of the mucus film to the typical length-scale of the sinus. The coordinate *Y* is a local stretched coordinate normal to the boundary; *Y* = 0 denotes the cavity wall and *Y* = *H*_c_ denotes the free surface of the fluid. Note that the reference scales defined above correspond to the classical thin-film approximation over curved substrates [24, 29, 42]. Note also that we have purposely defined a generic velocity scale 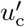, to show how the velocity scale emerges naturally from the equations governing fluid flow.

After the governing equations are rendered dimensionless using (A1), we identify the dominant balance in each equation by retaining only the leading order terms, i.e. the terms in each equation with the lowest powers of *ϵ*. This yields the continuity equation in the thin-film limit,

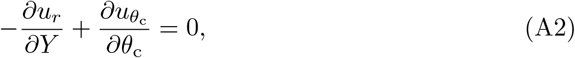

the dimensionless *r*^′^-momentum (or, *Y* -momentum) equation in the thin-film limit,

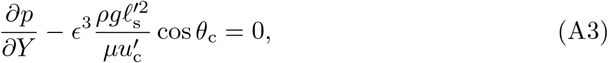

and the dimensionless *θ*_c_-momentum equation in the thin-film limit,

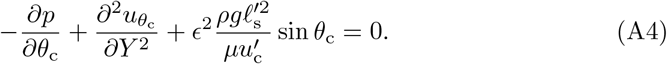

Eqn. (A4) provides the characteristic velocity scale,

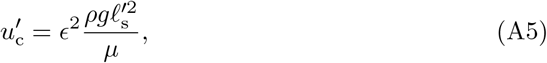

that we employ in all our derivations. Using this scale, eqns. (A3) and (A4) simplify to:

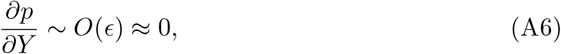

and,

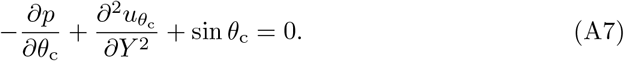

The system in Eqns. (A2), (A6) and (A7) is supplemented by: (i) boundary conditions (BCs) for the fluid velocity (*uθ*_c_, *u*_*r*_) at the walls of the circle, (ii) BCs for the fluid stress at the free surface of the thin film, and, (iii) a kinematic boundary condition relating the fluid’s velocity at the free surface to the film deformation. The first of these set of BCs is given by:

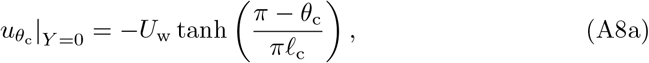

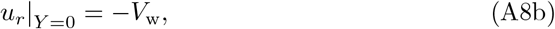

where,

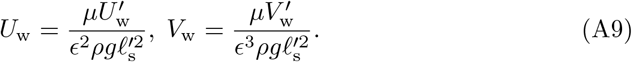

In the absence of surface tension gradients and any externally imposed stresses, the tangential stress in the fluid vanishes at the free surface:

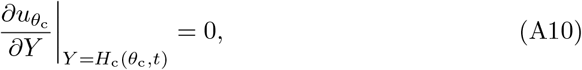

whereas the normal fluid stress undergoes a jump due to surface tension:

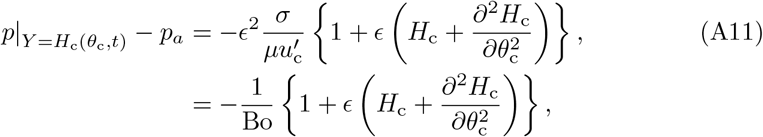

where *p*_*a*_ is the (uniform) air pressure in the cavity and *σ* is the surface tension of the air-fluid interface. In Eqn. (A11), the term within { } is the (in-plane) film curvature at the angular position *θ*_c_ as a function of the film thickness *H*_c_, and 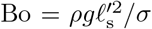 is the Bond number, which is a dimensionless measure of the relative importance of gravity and surface tension in driving the film. Finally, we have the kinematic boundary condition, relating the (leading order) fluid velocity at the free surface to the rate of deformation of the film:

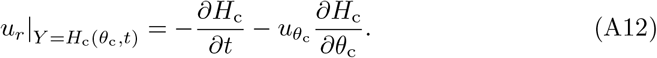

One can solve Eqn. (A7) subject to Eqn. (A8a), and Eqn. (A10) to obtain the following expression for the tangential fluid velocity:

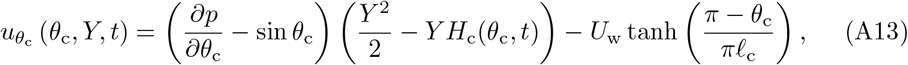

where the pressure gradient *∂p/∂θ*_c_ can be calculated using eqns. (A6) and (A11) as:

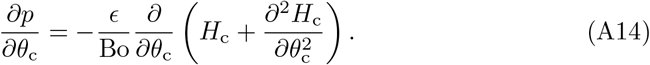

We can integrate Eqn. (A2) from *Y* = 0 to *Y* = *H*_c_ (*θ*_c_, *t*), and use eqns. (A8b), (A12), (A13), and the Leibniz integration rule:

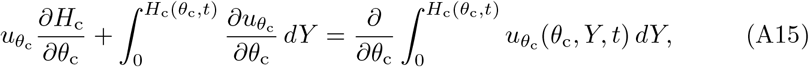

to arrive at the final thin film equation for circular geometry given in the main text’s eqn. (4),

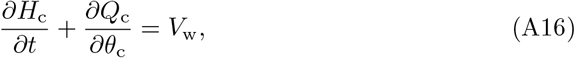

where,

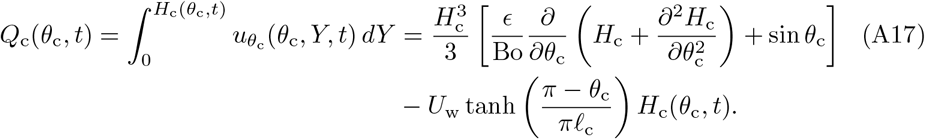

#### A.2 Description of the numerical method

We solve eqns. (A16) and (A17) numerically using a semi-implicit finite-difference scheme. We first expand Eqn. (A16) and write it as:

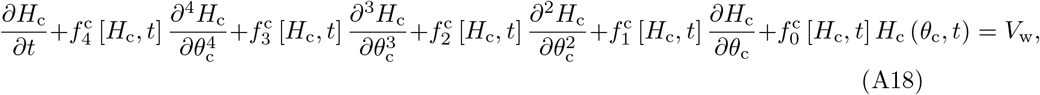

where,

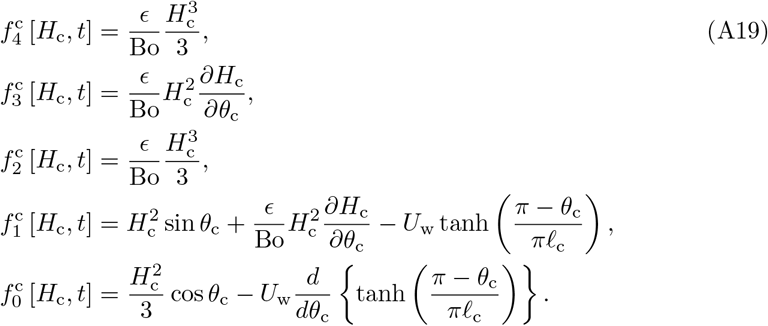

Due to geometric symmetry (see Fig. 2(a)), Eqn. (A18) is solved over the half-domain 0 ≤ *θ*_c_ ≤ *π*. We discretise Eqn. (A18) in space (i.e. the *θ*_c_ derivatives) using secondorder accurate finite difference approximations, and in time using an explicit Euler discretisation. The spatial discretisations are backward- and forward-biased at *θ*_c_ = 0 and *θ*_c_ = *π*, respectively. To obtain *H*_c_ (*θ*_c_, *t*_*n*+1_), the functions *f*_*i*_ [*H*_c_, *t*] are evaluated at the (previous) time-step *t*_*n*_, whereas the derivatives of *H*_c_ are evaluated at the desired/present time-step *t*_*n*+1_. The numerical simulations are initialised by prescribing a uniform initial thickness *H*_c_ (*θ*_c_, *t* = 0) = 0.1.

#### A.3 Validation of the numerical method

We validate our numerical implementation in the limit *U*_w_ = *V*_w_ = 0, by comparing our results to existing solutions for the height of a thin film draining on the outer surface of a cylinder [27]. We emphasise that this comparison can be made because the governing equation for our problem (where the film can be thought as developing inside a cylinder) is exactly the same as the problem where the film develops outside the cylinder. This is true even if gravity acts in opposite directions (with respect to the substrate normal extending into the fluid) depending on whether the film develops inside or outside the cylinder. The reason being, in the thin-film limit, the influence of gravity normal to the substrate is generally sub-dominant to leading order in 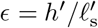 (see eqns. (A3), (A5) and (A6)) [45, 46]. The time evolution of a thin film draining passively (under the influence of gravity) outside/inside a cylinder–for a specific Bond number–is shown in Fig. A2a, and the agreement between our results and those of Ref. [27] validates our numerical method.

In addition to validating our numerical scheme in a limiting case, we confirm the resolution independence of the results provided in the main text. Towards this, we numerically solve Eqn. (A18) for increasing resolutions *N*_*θ*_ (i.e. the number of discretised points at which the film height *H*_c_ is computed) and notice negligible change in the steady-state solution; some examples are provided in Fig. A2b.

## Appendix B 3D/spherical system

### B.1 Derivation of the thin-film equation

For the spherical geometry, we work in spherical coordinates (*r*^′^, *θ*_s_, *ϕ*). Since we are interested in an axisymmetric flow, the *ϕ*-component of the velocity and all derivatives with respect to the *ϕ*-coordinate are identically zero, i.e. 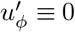 and *∂*()*/∂ϕ* ≡ 0. The derivation of the thin-film equation for the spherical geometry follows similar steps as that discussed for the circular case, and we provide here the main (dimensionless) equations required for deriving Eqn. (7). The continuity equation is given by,

**Fig. A2:**
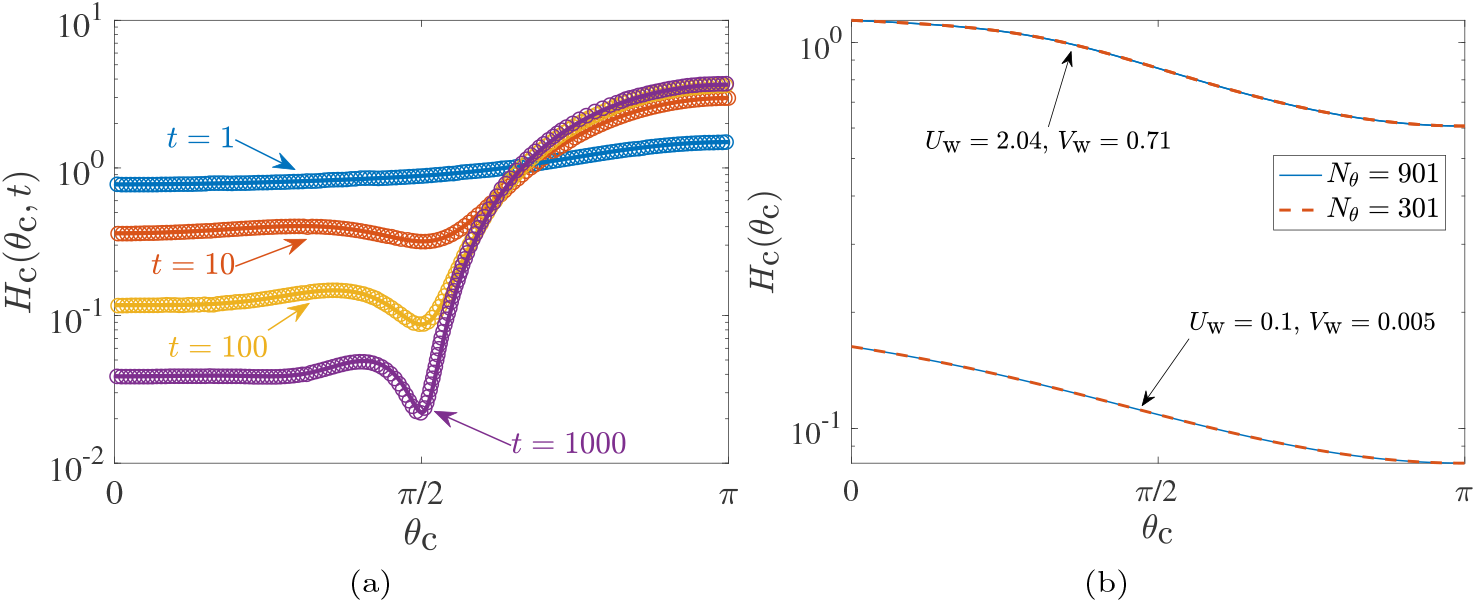
(a) Comparison between the finite-difference solution in the 2D case, Eqn. (A16) (with *U*_w_ = *V*_w_ = 0) in the present work and the numerical solution of Ref. [27] (see their Fig. 2) for the drainage of a thin film over a circular cylinder, at dimensionless times *t* = 1, 10, 100, 1000. The symbols denote numerical solutions of Ref. [27] and the solid lines denote our solutions. The Bond number for these film profiles is such that *ϵ/*Bo = (*π*^2^ − 8)*/*(8*π*) ≈ 0.07439. (b) Convergence of the numerical solution to Eqn. (A18) for different numbers of grid points *N*_*θ*_, shown for two different values of (*U*_w_, *V*_w_).

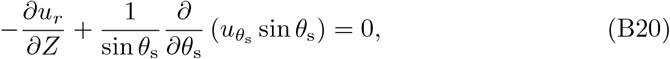

the *r*^′^-momentum equation is,

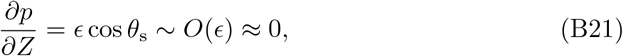

where *Z*, just like *Y* in the circular case, is a local stretched coordinate normal to the walls of the spherical cavity, such that *Z* = 0 denotes the cavity walls and *Z* = *H*_s_ denotes the free surface of the fluid (Fig. A1b). The *θ*_s_-momentum equation is given by,

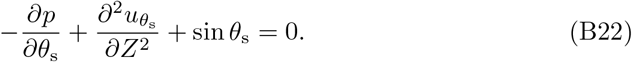

The boundary conditions for the velocities (*uθ*_s_, *u*_*r*_) at *Z* = 0 are, just as in the circular case:

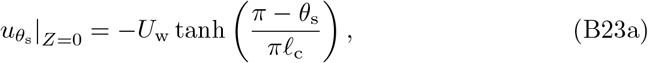

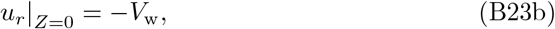

where, (*U*_w_, *V*_w_) were defined before in Eqn. (A9).

The tangential stress boundary condition is:

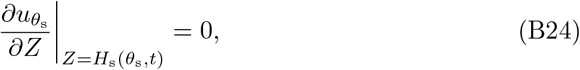

whereas the normal fluid stress condition is:

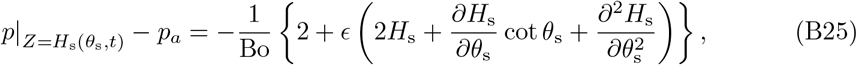

where, again, the term within { } is the film curvature at the angular position *θ*_s_ as a function of the film thickness *H*_s_. We also have the kinematic boundary condition as:

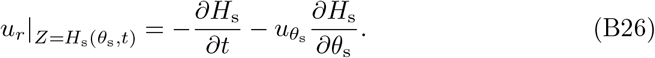

Following exactly the same steps as in the circular case, we obtain the following expression for the fluid velocity,

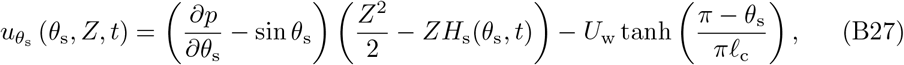

where, now the pressure gradient is derived from Eqn. (B25) as,

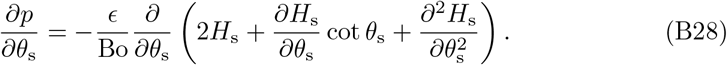

Integrating Eqn. (B20) from *Z* = 0 to *Z* = *H*_s_ (*θ*_s_, *t*), and using eqns. (B23b), (B26), (B27), and the Leibniz integration rule:

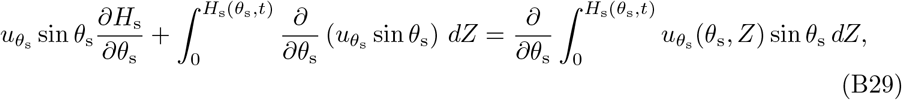

we obtain the final thin film equation for spherical geometry given in the main text,

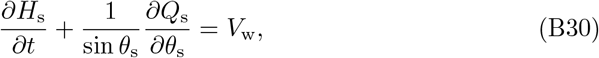

where,

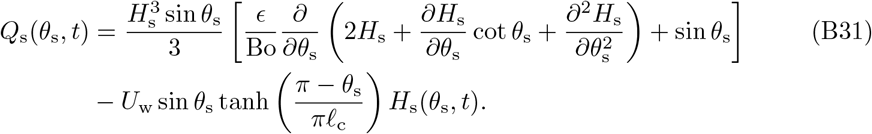

#### B.2 Details of the numerical method

The numerical method is identical to that used for circular geometry, except that the domain extends from 0 *< θ*_e_ ≤ *θ*_s_ ≤ *π* (see Fig. 2(b)). We do not repeat the details of the numerical method and provide here just the expanded form of Eqn. (B30):

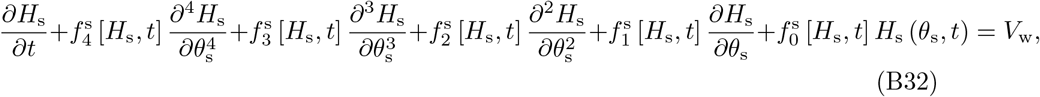

where,

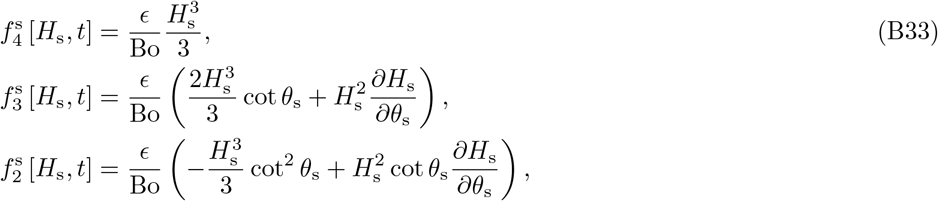

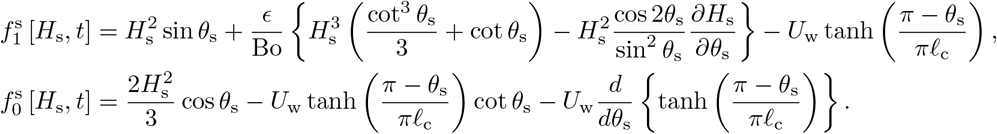

#### B.3 Validation of the numerical method

We validate our numerical solution in the limit *U*_w_ = *V*_w_ = 0 and *θ*_e_ = 0, by comparing our solution to existing results for the height of a thin film draining on the outer surface of a sphere [26]. As in the 2D case, it is important to note that this comparison is possible because, in the thin-film limit, the governing equation for our problem (where the film develops inside a sphere) is the same as the problem where the film develops outside the sphere. The comparison–for a prescribed Bond number–is shown in Fig. B3a, and the agreement between our results and those of Ref. [26] validates our numerical method. The convergence of the solution to Eqn. (B32) with respect to the resolution of spatial discretization (i.e. the number of points *N*_*θ*_ at which *H*_s_ is computed), is shown in Fig. B3b.

**Fig. B3:**
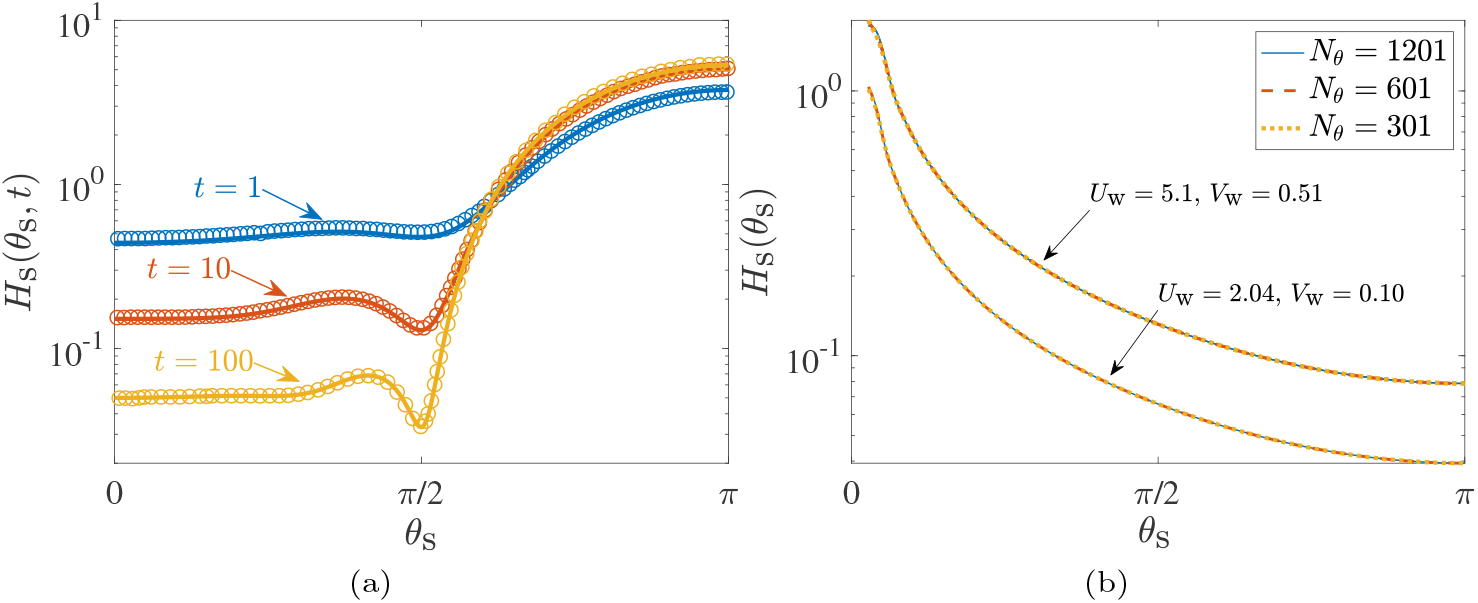
Comparison between the finite-difference 3D solution to Eqn. (B30) (with *U*_w_ = *V*_w_ = 0) in the present work and the numerical solution of Ref. [26] (see their Fig. 2) for the drainage of a thin film over a sphere, at dimensionless times *t* = 1, 10, 100. The symbols denote numerical solutions of Ref. [26] and the solid lines denote our solutions. The Bond number for these film profiles is such that *ϵ/*Bo = 1*/*24. (b) Convergence of the numerical solution to Eqn. (B32) for different numbers of discretization grid points *N*_*θ*_, shown for two different values of (*U*_w_, *V*_w_).

